# Relating network analyses to phylogenetic relatedness to infer protistan co-occurrences and co-exclusions in marine and terrestrial environments

**DOI:** 10.1101/2020.04.27.063685

**Authors:** Guillaume Lentendu, Micah Dunthorn

## Abstract

We used two large-scale metabarcoding datasets to evaluate phylogenetic signals at global marine and regional terrestrial scales using co-occurrence and co-exclusion networks. Phylogenetic relatedness was estimated using either global pairwise sequence distance or phylogenetic distance and the significance of observed patterns relating networks and phylogenies were evaluated against two null models. In all datasets, we found that phylogenetically close OTUs significantly co-occurred more often, and OTUs with intermediate phylogenetic relatedness co-occurred less often, than expected by chance. Phylogenetically close OTUs co-excluded less often than expected by chance in the marine datasets only. Simultaneous excess of co-occurrences and co-exclusions were observed in the inversion zone between close and intermediate phylogenetic distance classes in marine surface. Similar patterns were observed by using either pairwise sequence or phylogenetic distances, and by using both null models. These results suggest that environmental filtering and dispersal limitation are the preponderant forces driving co-occurrence of protists in both environments, while signal of competitive exclusion was only detected in the marine surface environment. The discrepancy in the co-exclusion pattern is potentially linked to the individual environments: water bodies are more homogeneous while tropical forest soils contain a myriad of nutrient rich micro-environment reducing the strength of mutual exclusion.

## Introduction

There is a long history of research trying to elucidate why species are present in a specific environment and why multiple species are found together (Darwin, 1859; Gause, 1934; Humboldt & Bonpland, 1805). Species sharing the same ecological niche tend to co-occur due to environmental filtering and dispersal limitation. In turn, closely-related species are more likely to co-occur due to their shared evolutionary history (e.g., common ancestor, shared traits) and their potential limited dispersal- and establishment-abilities. These processes can be balanced by densitydependent negative biotic interactions, like competitive exclusion when functionally similar species are after the same resource and co-exclude themselves. Environmental filtering and dispersal limitation have been identified as the main drivers shaping the assembly of most protists in the environment (Boenigk et al., 2018; de Vargas et al., 2015; del Campo et al., 2015; Lentendu et al., 2018; Mahé et al., 2017; Singer et al., 2018; Wetzel et al., 2012), while competition have been only formally tested in laboratory conditions (Saleem, Fetzer, Dormann, Harms, & Chatzinotas, 2012; Violle, Nemergut, Pu, & Jiang, 2011). These mechanisms have been largely evaluated for macroorganisms in different environments (Cavender-Bares, Kozak, Fine, & Kembel, 2009; Kraft et al., 2015), but have not yet been broadly evaluated for microbes in natural environments, for which community ecological analyses have rarely integrated phylogenetic information.

In environmental microbial ecology, environmental filtering is often considered as the prevalent limiting parameter of species occurrence (Khomich, Kauserud, Logares, Rasconi, & Andersen, 2017; Lauber, Strickland, Bradford, & Fierer, 2008; Lentendu et al., 2018; Philippot et al., 2010; Singer et al., 2018; Tedersoo et al., 2016; Weißbecker et al., 2018; Zinger et al., 2011) and is directly linked to the ecological niche of microbes (i.e., the set of abiotic parameter ranges in which a species can leave in). Ecological niche of microbes is hardly measurable without cultivation (Lennon, Aanderud, Lehmkuhl, & Schoolmaster, 2012; J. B. H. Martiny, Jones, Lennon, & Martiny, 2015), so that for large scale studies mostly based on non-cultivable microbes, function and functional similarity are either deducted from taxonomic or phylogenetic similarity of recovered sequences. Environmental filtering is inferred from the non-random co-occurrence of members of a taxa or a clade or from clade or taxa occurring in a restricted set of habitats. Thus, environmental filtering, when analyses in a phylogenetic context, often assumes phylogenetic niche conservatism, that is the long-term retention of ecological traits among closely related species (Wiens et al., 2010). Phylogenetic niche conservatism was shown in bacteria, mainly for complex functional traits which are conserved inside single clades (A. C. Martiny et al., 2013). Under phylogenetic niche conservatism, evolutionary close species are more likely to share the same ecological niche and thus tend to be filtered into the same habitats. With this assumption, environmental filtering can be tested using measures of phylogenetic divergence (e.g. MPD, MNTD, but see Tucker et al., 2017), with phylogenetic over-clustering (i.e. low phylogenetic divergence) being interpreted as sign for environmental filtering. This sample-wide approach has been used to support environmental filtering of trees, bacteria and protists along habitat and nutrient gradients (Horner-Devine & Bohannan, 2006; Kembel & Hubbell, 2006; Singer et al., 2018). However, it appears that most studies concluding on environmental filtering do not account for biotic interactions which could produce similar results (Kraft et al., 2015).

Competition is long known experimentally and it was hypothesized to drive co-exclusion in an initial experimental study involving protists (Gause, 1934). Competitive exclusion was first viewed as an evolutionary pressure which trigger trait divergence of related species, allowing them to escape competition and to persist in the same habitat, as originally observed for Darwin’s finches (Darwin, 1859). This assumption was further formalized with the phylogenetic limiting similarity hypothesis, in which phylogenetic related species do compete stronger due to niche overlap thus limiting the number of related species which can coexist (Macarthur & Levins, 1967). By assuming phylogenetic niche conservatism, it is expected that competitive exclusion will only affect closely related species, so that phylogenetic over-dispersion (i.e. high phylogenetic divergence) of natural communities is interpreted as a sign of competitive exclusion. This approach have allowed to identify one tree family presenting signs of competitive exclusion in a tropical forest (Manel et al., 2014). However, competition do not necessarily lead to exclusion when for example competition is symmetric or when other biotic interactions (e.g., mutualism or herbivory) reduce or neutralize the competition (Lamb & Cahill Jr., 2008; Müller, Hauzy, & Hulot, 2012; Olff & Ritchie, 1998). Further experimental evidences have shown that for protists species, competition will more quickly lead to exclusion when species are phylogenetically related, with a direct relation to phylogenetically conserved traits (e.g. mouth size Violle et al., 2011). The “paradox of the plankton” was also considered to be an opposite example of competitive exclusion, with the coexistence of high number of species using the same resources (Hutchinson, 1961). It was however shown that this pattern is explained by the competition itself which only leads to short term exclusion in a system never reaching an equilibrium (Huisman & Weissing, 1999). In plant ecology, studies measuring competition strength have shown that depending on clades or depending on soil conditions, there will be more or less competition between related species, so that no generalization of the ‘competition-relatedness’ hypothesis is possible (Burns & Strauss, 2011; Cahill, Kembel, Lamb, & Keddy, 2008). The exclusion of closely related species due to competition can thus be viewed as a special case of the coexistence theory (Mayfield & Levine, 2010). But so far, no large-scale study has tested for phylogenetic overdispersion and exclusion patterns in protists.

In today’s very large environmental sequencing datasets, microbial taxa are characterize using operational taxonomic units (OTU) which are used as proxy to molecular species (Blaxter et al., 2005). At the same time, co-occurrence and co-exclusion networks analyses have become standard in environmental microbial ecology, with a predominance of studies interested in cooccurrence patterns among and between taxonomic groups with a presumed function (Chow, Kim, Sachdeva, Caron, & Fuhrman, 2014; Lima-Mendez et al., 2015; Milici et al., 2016; Steele et al., 2011). To contrast with phylogenetic divergence analyses conducted at the sample level, cooccurrence and co-exclusion network analyses allow to extract statistically significant pair of co-occurring/co-excluding OTUs at the whole study level. By comparing observed co-occurrences to random co-occurrences among the regional pool of OTUs, signal for potential biotic interactions like parasitism, predation or viral infection have been disclosed (Lentendu et al., 2014; Lima-Mendez et al., 2015; Steele et al., 2011). By taking advantage of the modularity structure of the cooccurrence networks, microbial occurrences have also been linked to habitat preference, which can be interpreted as the signal for environmental filtering (de Menezes et al., 2014; Lentendu et al., 2014; Milici et al., 2016; Morriën et al., 2017). However, studies have yet to integrate the phylogenetic relatedness as an explaining parameter for network structure.

Here we describe a new analytical approach that aims to evaluate community assembly processes by decomposing the co-occurrence and co-exclusion networks among phylogenetic relatedness classes. By looking at excess or deficit of co-occurrence or co-exclusion in class of organism with increasing phylogenetic relatedness, we can test the possible assembly mechanisms in natural protistan communities. Under the assumption of phylogenetic niche conservatism, we tested the following hypotheses: i) if environmental filtering dominate, phylogenetically related OTUs will co-occur more and co-exclude less often than expected by chance and conversely for pairs of OTUs with intermediate phylogenetic relatedness; ii) if competitive exclusion dominate, phylogenetically related OTUs will co-occur less and co-exclude more often than expected by chance and conversely for pairs of OTUs with intermediate phylogenetic relatedness. To evaluate these hypotheses, we use two of the largest environmental sequencing protist datasets to date: the global marine subsurface dataset of de Vargas et al. (de Vargas et al., 2015), and the Neotropical rainforest soil dataset of Mahé et al. (2017). While both studies were primarily concerned by describing the occurrence of different taxa in different water bodies or forest soils, the current study try to evaluate how phylogenetic relatedness could explain the distributions of protists at global and regional scales.

## Material and Methods

The complete bash and R (R Core Team, 2017) scripts to reproduce the analyses are provided in HTML format (**File S1**). The full network calculation procedure is also available as a stand-alone software with multiple matrix normalization, randomization and thresholding options (https://github.com/lentendu/NetworkNullHPC).

### Datasets

Two large-scale environmental sequencing projects that focused on protistan diversity were used here (available upon request). Protistan OTUs from the world’s open oceans and seas came from de Vargas et al. (2015). This marine dataset is composed of 355 samples collected at the surface and deep chlorophyll maximum (DCM), which produced 366,800,845 protist reads of the V9 hypervariable region of the SSU-rRNA locus that clustered into 302,663 OTUs. To allow for comparison, the version of this marine dataset used here was re-analyzed by Mahé et al. (2017). All filter-size classes libraries of either the surface or DCM at a single station were pooled together, thus the number of samples used here reduced to 47 for surface and 32 for DCM waters. Protistan OTUs from three lowland Neotropical rainforests came from Mahé et al. (2017). This terrestrial dataset is composed of 144 samples collected at the soil surface, which produced 46,652,206 protist reads of the V4 hyper-variable region of the SSU-rRNA locus that clustered into 26,860 OTUs. For sampling and sequencing information see the original publications (de Vargas et al., 2015; Mahé et al., 2017); for bioinformatic pipeline of reads cleaning, clustering with Swarm v2 (Mahé, Rognes, Quince, de Vargas, & Dunthorn, 2015), and taxonomic assignments using the Protist Ribosomal Reference database (Guillou et al., 2013) to protists see Mahé et al. (2017). It is important to note that this reference database does not reflect the exact current international agreement on the taxonomy of protists (S. M. Adl et al., 2019) and each taxonomic path is reduced to eight taxonomic levels.

### Co-occurrence and co-exclusion networks

To infer protistan co-occurrences and co-exclusions from the marine and terrestrial datasets, networks were constructed using OTUs following Connor et al. (2017). This method infer positive correlations (co-occurrences), which was expanded here to also infer negative correlations (coexclusions). Resulting networks were composed of nodes (OTUs) that were connected by edges to one or more other nodes; these edges were either instances of co-occurrences or co-exclusions. First, to reduce computational load, OTUs occurring in less than 30% of marine and 10% of terrestrial samples were removed as well as samples with less than 20% of median read counts per sample in the terrestrial dataset. Low occurrence OTUs would never show any significant cooccurrence or co-exclusion using this method (Connor et al., 2017). The OTUs which passed the occurrence filter are later referred as the candidate OTUs. Second, read counts per sample were normalized using the log-ratio count method: reads were log transformed in order to reduce abundance bias due to PCR; counts were then normalized per sample to a median sequencing depth by multiplying read counts by the ratio of a minimum expected sequencing depth (half the median of original sample’s read count) by the sample’s total sum of read counts and rounding to integer. This normalization is preferable to rarefaction and/or relative abundance normalization, because it avoids random subsampling and variance inflation while taking into account the compositionality of the data (Gloor, Macklaim, Pawlowsky-Glahn, & Egozcue, 2017; McMurdie & Holmes, 2014). Third, random noise was added to the normalized matrices in order to break ties when calculating Spearman’s rank correlation coefficient (*rho*). Fourth, this random noise addition was repeated 1000 times (i.e., Monte Carlo sampling) to obtain a normal distribution of Spearman’s *rho*. Fifth, the thresholds to detect a biological significant positive (co-occurrence) or negative (co-exclusion) correlation were determined with randomly shuffled and noise-added OTU matrices. This threshold was set at the Spearman’s *rho* for which the largest connected component of a network, build with edges equal and above this threshold for co-occurrence, or equal and below this threshold for co-exclusion, contains less than 1% of the total OTU number in at least 90 % of the 1000 random OTU matrices. The random shuffling was based on OTU abundance swaps constrained to each sample and was prefer to the original full count shuffling without fixed row and column sums because it preserved the slight positive shift in Spearman’s rho as observed in natural communities (**Figure S1**). Sixth, observed edges with a Spearman’s *rho* above or below the selected threshold in at least 90 % of the Monte Carlo sampling and with corrected Spearman’s *rho* p.values (Benjamini & Hochberg, 1995) ≤ 0.01 in at least 90 % of the Monte Carlo sampling were considered as biological co-occurrence or co-exclusion, respectively. This procedure sets Spearman’s *rho* co-occurrence thresholds at 0.58 for marine surface, 0.68 for marine DCM, and 0.45 for terrestrial. Spearman’s *rho* co-exclusion thresholds were set at −0.52 for marine surface and −0.64 for marine DCM, and −0.24 for terrestrial (**Table 1**).

**Table 1.** Network parameters

| network | dataset | samples | candidate<br>OTUs* | reads | Spearman's<br>*rho*<br>threshold | network<br>OTUs<br>§ | network<br>reads | candidate<br>correlations<br>§ | significant<br>correlations<br>& | average<br>network<br>degree | average network<br>path length |
| --- | --- | --- | --- | --- | --- | --- | --- | --- | --- | --- | --- |
| co-occurrence | marine<br>surface | 47 | 8274 | 1.1e+8 | 0.58 | 4351<br>(52.6 %) | 8.2e+7 | 3.4e+7 | 49616<br>(0.14 %) | 22.8 | 4.7 |
|  | marine<br>DCM | 32 | 10760 | 6.5e+7 | 0.68 | 3575<br>(33.2 %) | 3.6e+7 | 5.8e+7 | 25306<br>(0.04 %) | 14.2 | 6.2 |
|  | Neotropical<br>soil | 114 | 687 | 1.8e+7 | 0.45 | 83<br>(12.1 %) | 5.2e+6 | 2.4e+5 | 373<br>(0.16 %) | 9.0 | 2.2 |
| co-exclusion | marine<br>surface | 47 | 8274 | 1.1e+8 | -0.52 | 4265<br>(51.5 %) | 8.1e+7 | 3.4e+7 | 29873<br>(0.09 %) | 14.0 | 4.0 |
|  | marine<br>DCM | 32 | 10760 | 6.5e+7 | -0.64 | 3478<br>(32.3 %) | 3.5e+7 | 5.8e+7 | 13760<br>(0.02 %) | 7.9 | 5.1 |
|  | Neotropical<br>soil | 114 | 687 | 1.8e+7 | -0.24 | 41<br>(6 %) | 4.8e+6 | 2.4e+5 | 54<br>(0.02 %) | 2.6 | 3.4 |

\* OTU of the original dataset occurring in at least 30 % of marine or 10 % of terrestrial samples
§ percentage of candidate OTUs in brackets
§ total number of potential edges between network OTUs
& number of edges in the network; percentage of potential edges in brackets

### Pairwise sequence and phylogenetic distances

To infer the phylogenetic relatedness between the OTUs (nodes) in the constructed co-occurrence or co-exclusion networks, the OTU representatives (the most abundant strictly-identical amplicon) were used. These phylogenetic relatedness values between the OTUs were then overlaid along the edges in the networks. Two methods were used to infer the phylogenetic relatedness. First, pairwise sequence distances were calculated using a Needleman-Wunsh approximation as implemented in SUMATRA v1.0.34 (Mercier, Boyer, Bonin, & Coissac, 2013). This global pairwise sequence comparison did not account for any model of evolution. Second, phylogenetic distances were calculated by aligning the sequences using the FFT-NS-i strategy in MAFFT v7.407 (Katoh & Standley, 2013) and by finding the best maximum-likelihood tree using the GTRCAT model in RAxML 8.2.12 (Stamatakis, 2014) with 256 random starting trees. The phylogenetic distance between each tree tip was then calculated with the “cophenetic” function in R (R Core Team, 2017).

### Null models

To infer if the associations between the networks (both co-occurrences or co-exclusions) and the phylogenetic relatedness differed significantly from randomness, two null models were constructed. Null model 1 followed Hardy (model 1s, 2008) by generating random phylogenetic relatednesses values between nodes. These random values were made by a custom script that randomly shuffled the tip of the phylogenetic tree limited to the OTUs presented in the co-occurrence or co-exclusion networks. The same random re-ordering of OTUs was applied to both pairwise sequence and phylogenetic distance matrices (i.e. re-ordering row and column names) and the distance value for each co-occurring or co-excluding OTU pair was extracted. Null model 1 aimed to test whether cooccurring or co-excluding OTUs are more or less phylogenetically related than expected by chance. Null model 2 followed Chung and Lu (2002) by generating random edges between nodes. In these random networks, the total amount of edges remained the same as in the observed network, but the number of edges from an individual node was drawn from a probability distribution in which edge probability depends on the cumulative observed degree of the two nodes involved. This null Chung-Lu model produced networks with characteristics (e.g. modularity, diameter, clustering coefficient) more similar to natural networks compared to the most widely used null Erdős-Rényi model (Connor et al., 2017), and thus minimizes the number of parameters modified compared to the observed network. The random networks were made using the “sample_fitness” function in the R igraph package (Csardi & Nepusz, 2006). Null model 2 aimed to test whether phylogenetically related OTU co-occurred or co-excluded more or less than expected by chance.

### Statistical analyses

Null model constructions were repeated 1,000 times in order to test for statistic difference with the observed data. Phylogenetic relatedness was aggregated step-wisely, using a step of 0.01 for pairwise sequence distances and a step of 0.1 for phylogenetic distances. For each distance class, the number of co-occurring or co-excluding OTUs was accounted in the observed and random networks and a non-parametric p-values was calculated as the amount of time the observed number of co-occurrence or co-exclusion was higher or lower than in the null models. Differences between the observed networks and the null models were considered significant if the p-values were ≤ 0.05. Results were summarized for each distance class into standardized effect size (SES), calculated following Gotelli & McCabe (2002). By convention, a SES is considered as strong if it is ≥ 2.

## Results

### Networks coverage

In order to test for a phylogenetic signal between co-occurring and co-excluding OTUs with different phylogenetic relatedness, co-occurrence and co-exclusion networks were related to pairwise sequence and phylogenetic distances: edges of connected OTUs in the networks were labeled with the phylogenetic relatedness distances and the number of edges in each distance class were compared to two null models. The marine protist networks consisted of 32 to 53 % of candidate OTUs, while terrestrial protist networks included only 6 to 12 % of candidate OTUs (**Table 1**). The network OTUs occurred in at least 32 % of marine surface, 37 % of marine DCM or 17 % of terrestrial samples. The terrestrial co-exclusion network included the lowest amount of candidate OTUs (6 %) and candidate edges (0.02 %) compared to all the other networks. The occurrence patterns of network OTUs were slightly skewed toward OTUs occurring in the highest number of samples and thus in the highest number of geographical units, compared to candidate OTUs (**Figure S2**). Marine protist networks included mainly OTUs occurring in 6 to 8 sea and oceans, and most candidate OTUs occurring in only 4 to 5 of this geographical units were not included in the networks. Terrestrial protists networks included mostly OTUs occurring in 2 to 3 forests while candidate OTUs occurring in a single forest were largely absent from the networks. The taxonomic coverage of network OTUs remain unchanged in marine datasets compared to candidate OTUs (**Figure S3**). OTUs of the two clades with the lowest abundance in the terrestrial dataset, Dinophyta and Haptophyta, were not included in the networks as well as Chlorophyta OTUs in the co-occurrence network and MAST (Marine Stramenopiles, polyphiletic basal clade; Massana, Campo, Sieracki, Audic, & Logares, 2014) OTUs in the co-exclusion network.

### Phylogenetic signal in co-occurrences networks

Using null model 1 in which phylogenetic relatedness values were randomized along the edges of the networks, co-occurring OTUs from the marine datasets had positive SES that were significant and strong for low pairwise sequence distances <0.27 and phylogenetic distances <1.7, and OTUs from the terrestrial dataset had positive SES that were significant and strong for pairwise sequence distances <0.25 and phylogenetic distances <0.9 (**Figure 2**). Conversely, OTUs from the marine datasets had negative SES that were significant for intermediate and large pairwise sequence distances (0.27 to 0.5) and phylogenetic distances (2.1 to 4.3 and 6.3 to 9.5 for marine surface, 1.9 to 6.3 and 7.7 to 9.2 for marine DCM), and OTUs from the terrestrial dataset had negative SES for intermediate values that were significant in only four pairwise sequence distance classes (0.28 to 0.35) and seven phylogenetic distance classes (1.1 to 2.3). Interestingly, co-occurrence in Neotropical soils showed significant positive SES for OTUs pairs with large dissimilarities at one pairwise sequence and four phylogenetic distance classes.

**Figure 1.**
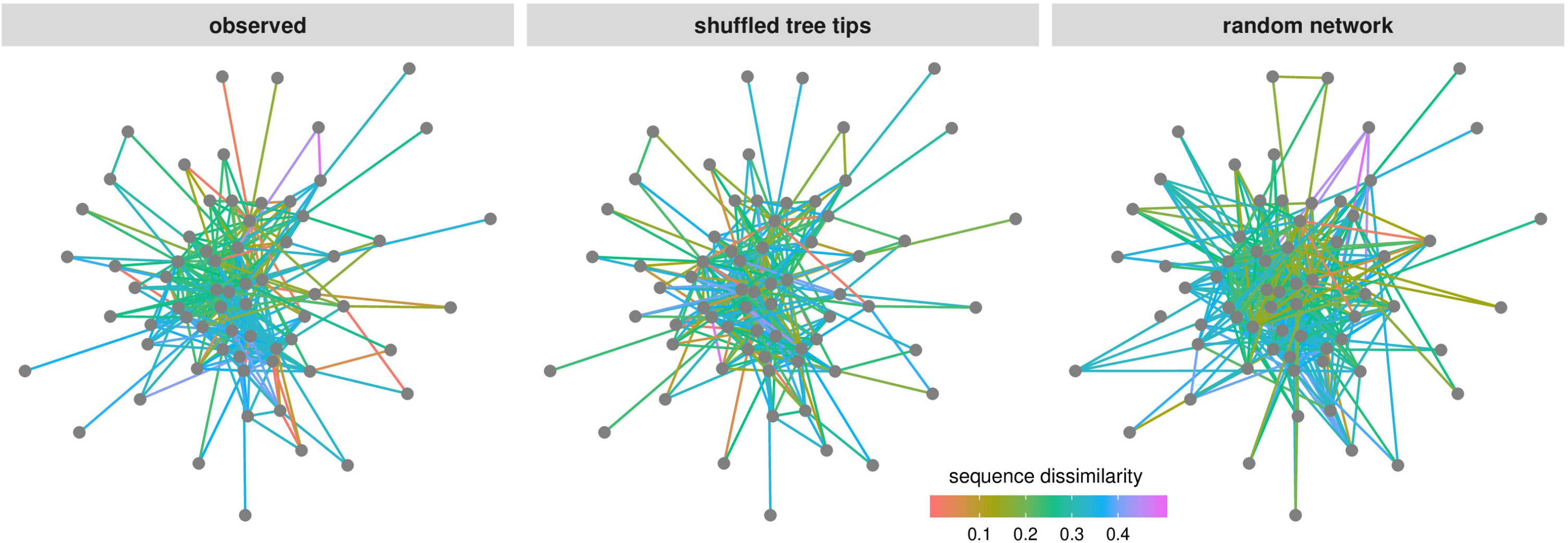
Null models effects on co-occurrence networks. Using the terrestrial protists co-occurrence network (observed) in which nodes are OTUs, edges are significant co-occurrences and edge colors are pairwise sequence dissimilarity. The first null model shuffle the pairwise sequence distance matrix (shuffled tree tips) while the second null model randomized the edges with a probability model (random network). The same approach was used for phylogenetic distance with phylogenetic tree tips shuffling in the first null model. The same computations were conducted on co-exclusion networks in which edges are significant co-exclusions.

**Figure 2.**
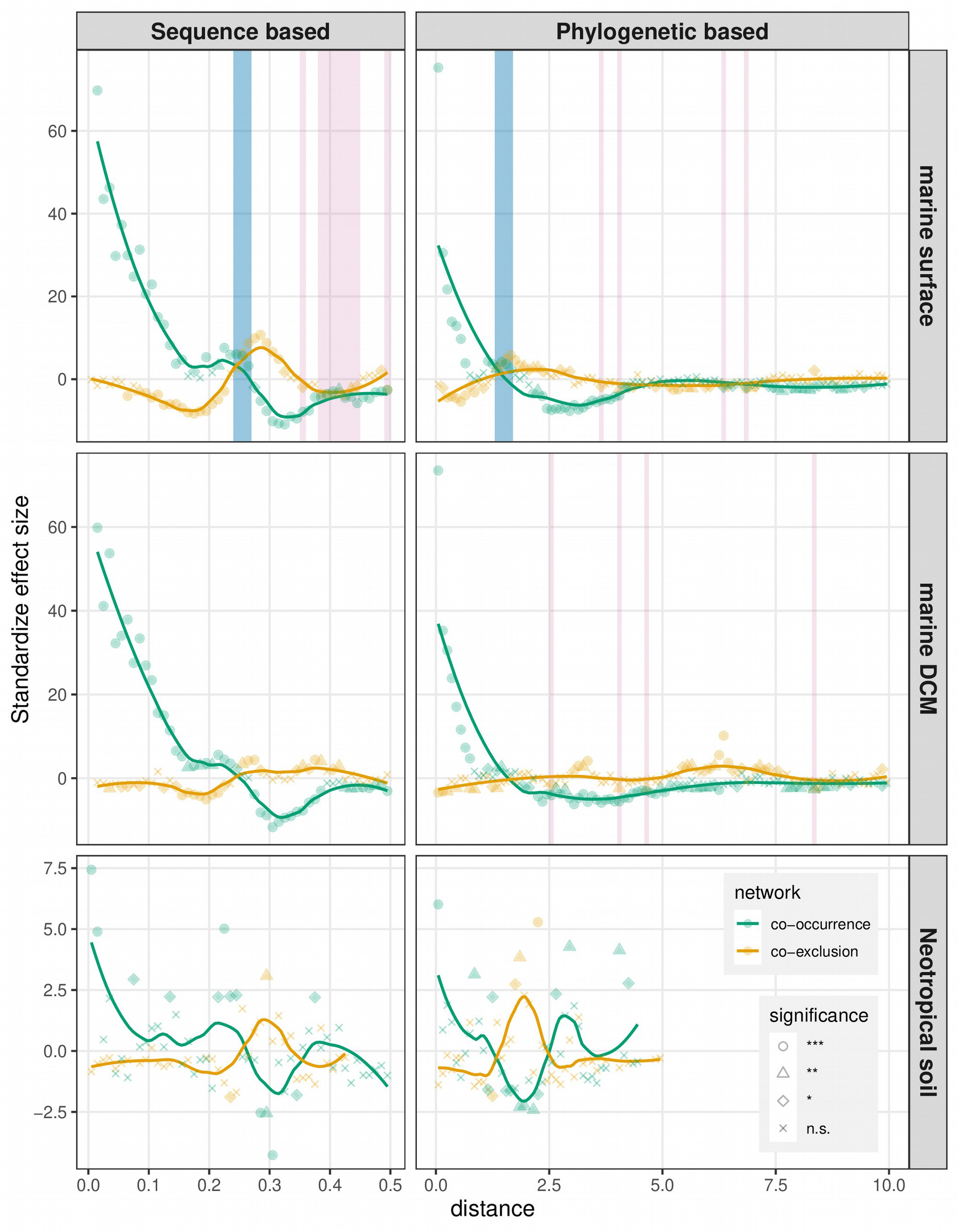
Standardize effect sizes (SES) in co-occurrence and co-exclusion networks compared to null models with shuffled phylogenetic tree tips (null model 1). SES were calculated separately for stepwise increased pairwise sequence genetic distances and phylogenetic distances. The number of OTU pairs connected by an edge in the observed networks was accounted for each distance class (from 0 to 0.5 with a 0.01 step for sequence based; from 0 to the maximum phylogenetic distance with a 0.1 step for phylogenetic based) and compared to the corresponding distance class reported from the randomized networks. Two-sided non-parametric p.values are inversely proportional to the amount of null models with a higher (for positive SES) or lower (for negative SES) amount of cooccurrence than in the observed network for each distance class. P.values below or equal to 0.05 were considered significant (* ≤ 0.05; ** ≤ 0.01; *** ≤ 0.001). Distance ranges highlighted in blue or red are for distances with excess (significant positive SES) or lack (significant negative SES) of edges in both co-occurrence and co-exclusion networks simultaneously.

Similar co-occurrence results to null model 1 were observed when using null model 2, in which the edges were randomized in the networks (**Figure S4**). Co-occurring OTUs from the marine datasets had positive SES that were significant and strong for pairwise sequence distances <0.23 and phylogenetic distances <1, and OTUs from the terrestrial dataset had positive SES that were significant and strong for pairwise sequence distances <0.04 and phylogenetic distances <0.9. And conversely, OTUs from the marine and terrestrial datasets had negative SES that were significant for intermediate pairwise sequence distances (>0.23) and phylogenetic distances (>1.1 to 6.7).

These results using the two null models mean that pairs of OTUs that are closely related phylogenetically co-occurred more often than expected by chance in the marine and terrestrial protistan communities, phylogenetically distant OTUs predominantly co-occured less often than expected by chance, and some phylogenetically far OTUs co-occurred more often than expected by chance. Additionally, for co-occurrences, using either pairwise sequence distances or phylogenetic distances in these comparisons results in similar SES values.

### Phylogenetic signal in co-exclusion networks

Using null model 1, co-excluding OTUs from the marine datasets had negative SES that were significant and strong for low pairwise sequence distances <0.23 and phylogenetic distances <1.1 in surface and <1.4 in DCM waters (**Figure 2**). Conversely, OTUs from the marine datasets had positive SES that were significant for intermediate pairwise sequence distances (surface: 0.24 to 0.33; DCM: 0.25 to 0.42) and phylogenetic distances (surface: 1.3 to 3; DCM: 3 to 3.4), while at higher distance classes a mix of significant positive and negative SES were retrieved. In the terrestrial dataset, however, no significant SES were observed except for the pairwise distance class between 0.29 an 0.3 and three phylogenetic distance classes between 1.7 and 2.3 with significant positive SES and pairwise distances between 0.23 and 0.24 and phylogenetic distances between 1.2 and 1.3 with a significant negative SES each.

Similar co-exclusion results to null model 1 were also observed when using null model 2 respectively (**Figure S4**). Co-excluding OTUs from the marine datasets had negative SES that were significant and strong for pairwise sequence distances <0.23 in surface and <0.16 in DCM waters, and phylogenetic distances <1.2 in surface <0.8 in DCM waters. No significant SES were observed in the terrestrial dataset except for the pairwise distance class between 0.29 an 0.3, three phylogenetic distance classes between 1.7 and 2.3 with significant positive SES and phylogenetic distances between 0 and 0.1 with a significant negative SES.

These results using the two null models mean that pairs of OTUs that are closely related phylogenetically co-excluded less often than expected by chance, and phylogenetically distant OTUs co-excluded more often than expected by chance, in the marine protistan communities. In the terrestrial protistan communities, though, there was an independence between phylogenetic relatedness and co-exclusion. Additionally, for co-exclusions, as in the co-occurrences, using either pairwise sequence distances or phylogenetic distances in these comparisons results in similar SES values.

### Synchrony and convergence in co-occurrence and co-exclusion patterns

In all datasets and for most distance classes, positive SES in co-occurrence networks were reflected by negative SES in co-exclusion networks and conversely. However, the negative SES in coexclusion networks for phylogenetically close OTUs were comparatively much lower or nonsignificant than the positive SES in the co-occurrence networks. These patterns are confirmed by the edge sampling along distance classes (**Figure S5**), with co-occurrence networks sampling most of candidate edges in low pairwise sequence and phylogenetic distances values, while co-exclusion networks lack of edges in those low distance values. It implied higher sampling of edges between OTUs from same genera in the marine datasets or from same species in the terrestrial dataset for cooccurrence networks (**Figure S6**).

For some distance classes there was, at the same time, significant positive or negative SES in both co-occurrence and co-exclusion networks (**Figure 2** and **S4**, shaded areas). This was particularly obvious for the marine surface dataset with null model 1 for which a SES inversion zone with positive SES in both co-occurrence and co-exclusion networks was observed over large ranges of pairwise sequence (0.24-0.27) and phylogenetic (1.3-1.7) distance classes (**Figure 2**). In the inversion zone, more than 80% of the co-occurrences and co-exclusions in the marine surface dataset were between taxa of different kingdoms and the distribution of edges among shared taxonomic levels did not differed significantly from the candidate edges in these same ranges (**Figure 3** and **S7**). A closer look at the taxonomic groups connected by co-occurrences and co-exclusions in the inversion zone revealed important shifts in proportion of edges compared to all candidate edges (**Figure S8**). Ciliophora were under-represented in both of these co-occurrence and co-exclusion sub-networks compared to all candidate edges as well as Apicomplexa, Bacillariophyta (diatoms), Dinophyta and Radiolaria in the co-occurrence sub-netwroks, while there were increase for almost all other pairs of clades in both sub-networks, in particular for Haptophyta in the co-occurrence sub-network and for Bacillariophyta vs. Dinophyta and Haptophyta in the co-exclusion sub-network (**Figure 4**). Interestingly, there were simultaneous excess of intra-clade cooccurrences and co-exclusions for Haptophyta, MAST and Telonemia and simultaneous lack of intra-class co-occurrences and co-exclusions for Ciliophora, Dinophyta and Radiolaria. The amount of changes was particularly important when comparing to the same sub-networks in the 0.24-027 pairwise sequence distance range of the marine DCM dataset (**Figure S9**). Edges involved less pairs of clades, and the lowest range of fold changes showed a much less divergent sampling of all potential edges than in the marine surface dataset, so that no inversion zone was visible for the marine DCM dataset.

**Figure 3.**
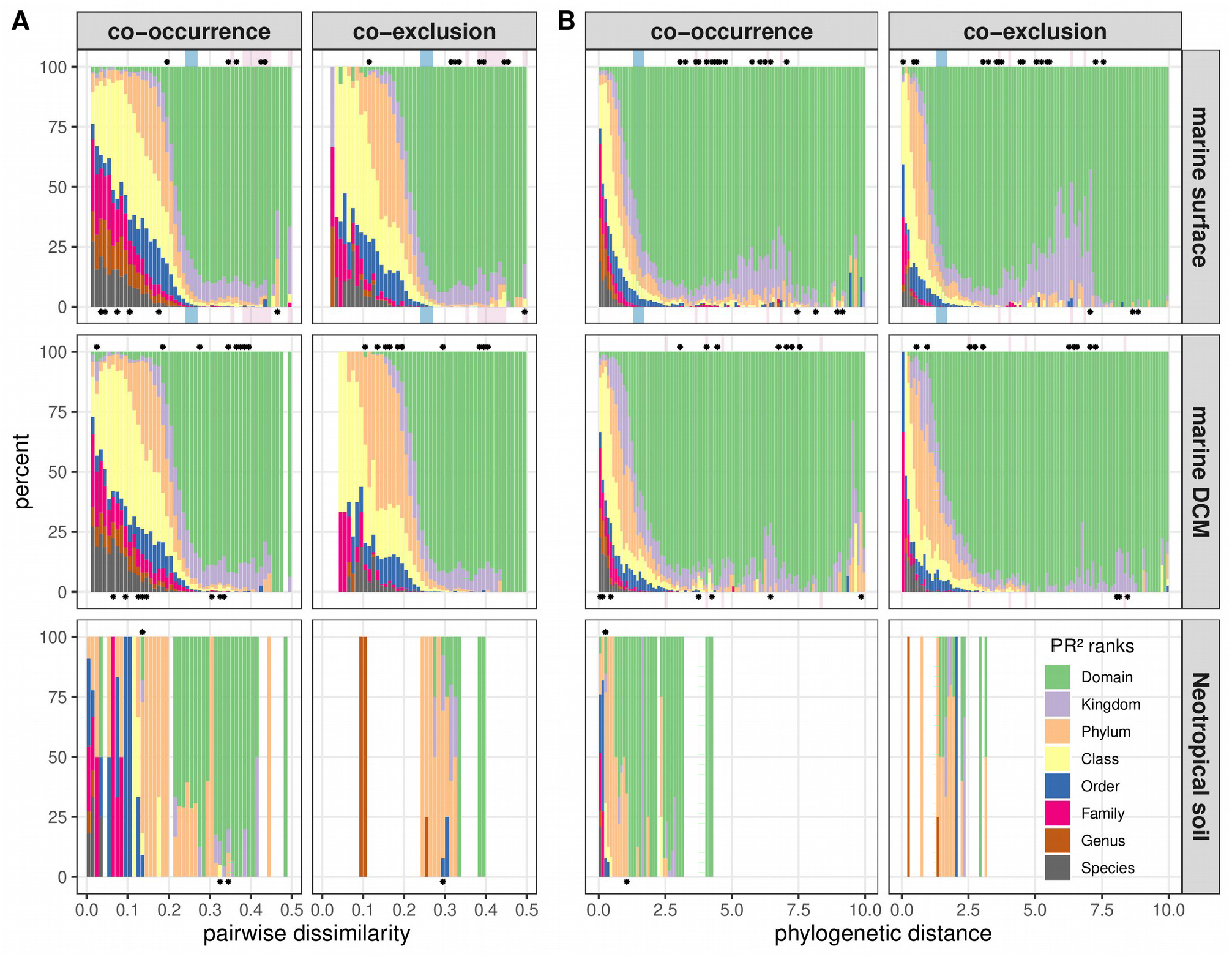
Distribution of taxonomic relationships between network connected OTUs for each pairwise sequence distance (a) and phylogenetic distance (b) classes. Blue and red shaded areas in the background are the distance classes with simultaneous positive or negative SES in both cooccurrence and co-exclusion networks using null model 1, as in Figure 2. Stars at the bottom of the bars indicate classes with significant deeper (toward species level) taxonomic ranks distribution compared to all candidate edges (Figure S7), stars at the top of the bars indicate classes with significant higher (toward domain level) taxonomic ranks distribution (Mann-Whitney test, p<0.05).

**Figure 4.**
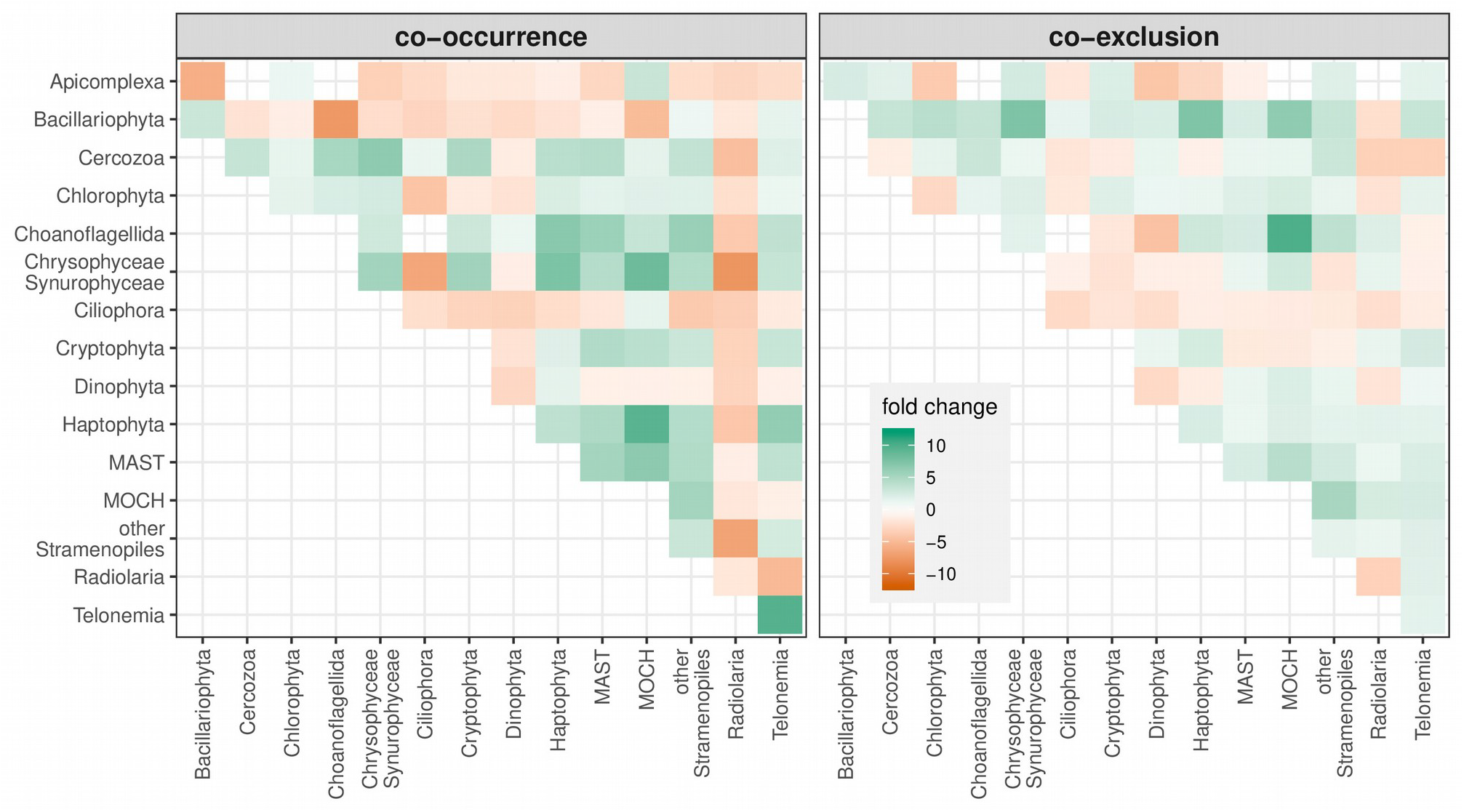
Fold changes in proportion of edges connecting the main clades in the marine surface dataset compared to all candidate edges in the pairwise sequence distance range of 0.24-0.27 (*i.e*. the largest range of distance with simultaneous positive SES in co-occurrence and co-exclusion networks when using the null model 1). The fold change color scale is identical to the one use for the marine DCM dataset (Figure S9).

## Discussion

We assessed the non-random phylogenetic relatedness of co-occurring and co-excluding OTUs in two of the largest environmental sequencing datasets of marine and terrestrial protists. By decomposing assembly patterns in phylogenetic relatedness classes and by comparing observed results to two null models, we could show that phylogenetic close OTUs co-occurred more often than expected by chance and that co-occurring OTUs are phylogenetically closer than expected by chance in both environments. The opposite trend was observed for OTUs with intermediate phylogenetic distances, which co-occurred less often than expected by chance. These co-occurrence results tend to support the preponderant effect of environmental filtering under the assumption of phylogenetic niche conservatism. These results could also be explained by the dispersal limitation of recently diverging taxa, which was demonstrated for the dominant protistan taxa in terrestrial dataset used here (Lentendu et al., 2018) or in marine ciliates (Azovsky, Chertoprud, Garlitska, Mazei, & Tikhonenkov, 2020).

Phylogenetic close OTUs were found to co-exclude less often than expected by chance while OTUs with intermediate phylogenetic distances co-excluded more often than expected by chance in the marine environments, in opposition to the co-occurrence patterns. There was, however, no clear limit between close and intermediate phylogenetic distances so that some distances classes displayed significant excess of both co-occurrences and co-exclusions in this transition zone in the marine surface dataset. In the terrestrial environment, however, co-exclusion was almost independent from phylogenetic relatedness. Under the assumption of phylogenetic niche conservatism, these co-exclusion patterns would also reflect the effect of environmental filtering in both marine surface and DCM waters, while neither environmental filtering nor competitive exclusion appeared to impact the distribution of protists in Neotropical soils. One explanation to this discrepancy would be the relatively higher level of homogenization and increased dispersal potential in the marine waters, which allows protists to more easily reach a suitable habitat, while the larger amount of soil protist microhabitats (M. S. Adl & Gupta, 2006) and the high local diversity in the Neotropics (Mahé et al., 2017) should blur the impact of environmental filtering and limit potential competitors to come into contact. Simultaneous excess of co-occurrences and coexclusions in Haptophyta and Telonemia in the SES inversion zone could reflect simultaneous effect of environmental filtering and competitive exclusion. While the “paradox of the plankton” and its resolution based on the theory of chaos support the co-occurrence of functionally similar plankton (Huisman & Weissing, 1999; Hutchinson, 1961), here we show that indeed phylogenetic related plankton co-occur but could simultaneously co-exclude themselves more than expected by chance at the marine surface. Other large-scale processes affect the assembly patterns of marine protist like the mean annual temperature responsible of the latitudinal diversity gradient or the sunlight exposure and currents responsible of the depth stratification in the water column (Giner et al., 2020; Ibarbalz et al., 2019). However, geographical structures, natural fluctuations and absence of equilibrium state in marine plankton communities are not enough to avoid exclusion among related organisms, as observed here, and would refute the existence of any plankton paradox under phylogenetic niche conservatism.

There are three novel aspects to this study. The first novel aspect was the use of null models to test the significance of phylogenetic relatedness structures in co-occurrence and co-exclusion networks. So far, only the relation between co-occurring/co-excluding protistan OTUs and their putative function or the change in network topology among habitats were tested in marine (Guidi et al., 2016; Lima-Mendez et al., 2015; Milici et al., 2016; Steele et al., 2011), freshwater (Debroas et al., 2017; Posch et al., 2015) and terrestrial environments (Lentendu et al., 2014; Ma et al., 2016; Xiong et al., 2017). In a network-based study on human microbiome combining analyses of phylogenetic relatedness and co-occurrence/co-exclusion networks, it was shown that co-occurrence between human bacterial OTUs were uniformly distributed among phylogenetic distances while coexclusions were mainly among phylogenetically distant OTUs (Faust et al., 2012). The lack of null model and/or statistical test on these observations, however, did not allow to determine whether biologic or random processes were responsible of the patterns. In a more recent study, global gut microbiome co-occurrence networks were found to have significant higher phylogenetic assortativity than in randomize networks overall (Tackmann, Matias Rodrigues, & von Mering, 2019), while size effect was not quantified at distinct distance classes and no interpretation was provided on these observations. Our new approach has the potential to uncover inter-dependencies between phylogenetic relatedness and co-occurrence and co-exclusion of any micro-organisms in any environment.

The second novel aspect is that we showed that both phylogenetic distance and pairwise sequence distance can both be used as measure of phylogenetic relatedness when applied to the analysis of protistan community assembly patterns. Previous protist studies used phylogenetic relatedness of protist to assess phylogenetic diversity based macroecological and biogeographical patterns (Bates et al., 2013; Lentendu et al., 2018; Singer et al., 2018), while pairwise sequence distances were only used during bioinformatic procedure for sequence clustering or sequence similarity networks (Forster et al., 2019; Mahé et al., 2015).

The third novel aspect was the decomposition of the co-occurrence and co-exclusion signals along phylogenetic distance classes. By using traditional index of phylogenetic divergence (e.g., net relatedness index), only one type of divergence could be assessed per sample or pair of samples, that is either clustering or overdispersion. By using the co-occurrence and co-exclusion patterns over all samples, here we investigated the multiple signals hold by communities over increasing phylogenetic distances for the whole analyzed regions. In the marine surface environment, at the SES inversion zone, both phylogenetic clustering and overdispersion take place at the same time. Independent to the origin of these patterns (competition could also lead to phylogenetic clustering, Mayfield & Levine, 2010), phylogenetic relatedness play a strong role in determining the assembly of marine and terrestrial protists.

There are three major assumptions to this study. The first major assumption was that there is phylogenetic niche conservatism between the OTUs (Wiens & Donoghue, 2004). This assumption allowed us to infer that phylogenetic close OTUs share more niche space than phylogenetically distant OTUs. This assumption allows us to interpret the significant excess in co-occurrence among phylogenetically close as a signal of environmental filtering and the absence of significant effect size in co-exclusion among phylogenetically close OTUs as signal for lack of environmental filtering and competitive exclusion. However, the assumption that evolutionary close OTUs share the same niche may not be true and it could be misleading to deduce pattern from process (Gerhold, Cahill, Winter, Bartish, & Prinzing, 2015). In such large dataset, there is a multitude of niche evolution scenarios which lead to the current distribution of protist in marine waters and Neotropical soils, and the apparent environmental filtering deducted here from the co-occurrence patterns could hide other processes at play which are not necessarily linked to phylogenetic niche conservatism. A modeling approach could also help to test for the reality of phylogenetic niche conservatism by protists (Münkemüller, Boucher, Thuiller, & Lavergne, 2015) but remains inapplicable for large datasets as analyzed here for which a large proportion of organisms are unknown (de Vargas et al., 2015; Mahé et al., 2017). Considering that current knowledge on traits and function is not sufficient to determine functional niche of most protists (Ramond et al., 2019), relating phylogeny to assembly patterns with the phylogenetic niche conservatism assumption is the most precise approach we can apply yet to find clues about large scale and whole community processes at play in protist community assembly.

The second major assumption to this study is that the OTUs are accurately estimating protistan species diversity. This assumption, which is made by most metabarcoding studies (Bik et al., 2012; Blaxter et al., 2005; Taberlet, Bonin, Zinger, & Coissac, 2018), allowed us to infer relative occurrences of each protist taxonomic unit among all samples of each datasets and allowed to infer the co-occurrence and co-exclusion networks. However, all clustering programs used to construct OTUs make assumptions about the best ways to handle the environmental sequencing data (Callahan et al., 2016; Caron & Hu, 2018; Mahé et al., 2015; Nebel, Pfabel, Stock, Dunthorn, & Stoeck, 2011; Rognes, Flouri, Nichols, Quince, & Mahé, 2016; Zhang, Kapli, Pavlidis, & Stamatakis, 2013) and these assumptions, along with the choice of molecular markers, may or may not lead to under- or over-estimations of species diversity. Here the reads were clustered into OTU with the program Swarm (Mahé et al., 2015; Mahé, Rognes, Quince, Vargas, & Dunthorn, 2014), which uses local clustering thresholds and a breaking phase to construct the OTUs. Swarm can partition the data into finer OTUs than programs that use global clustering thresholds, which may lead to over-splitting of species (Mahé et al., 2015); this over-splitting could potential explain the high positive SES in the smallest pairwise sequence and phylogenetic distance classes of cooccurrence networks.

The third assumption is that phylogenetic relatedness is correctly assessed with the analyzed genes. This assumption allowed us to infer strong interrelationship between phylogenetic distance and co-occurrence and co-exclusion patterns. The short and hyper-variable V4 and V9 fragments only provide partial phylogenetic signal of the full SSU-rRNA locus (Dunthorn et al., 2014), which is in-turn, only an approximation of the real protistan phylogenetic relatedness as assessed with whole genome sequencing (Burki, 2014). Besides, the genetic distances estimated between these two hyper-variable regions can be the same or drastically different depending on which taxa are being compared (Dunthorn, Klier, Bunge, & Stoeck, 2012; Hu et al., 2015; Tragin, Zingone, & Vaulot, 2018). The congruent results for protistan co-occurrences and co-exclusions derived from both pairwise sequence distances and phylogenetic distances shows that both type of distances can be used to infer phylogenetic relatedness. The congruent co-occurrence results for both global marine and Neotropical soil protists shows that both V4 and V9 fragments could deliver similar phylogenetic related assembly structure so that could be equally applied for large scale datasets.

By demonstrating the strong phylogenetic signals in co-occurrence and co-exclusion patterns of protists, we showed that global and regional assembly mechanisms are directly related to phylogenetic relatedness and are dominated by environmental filtering. We could not conclude that the simultaneous excess of co-occurrence and co-exclusion of phylogenetic related OTUs in the SES inversion zone of the marine surface communities is the result of intra-clade competitive exclusion, but we could only suspect it. Indeed, multiple other processes could lead to such pattern, like facilitation of phylogenetically distant species (Cahill et al., 2008; Gerhold et al., 2015; Kraft, Cornwell, Webb, & Ackerly, 2007). The co-exclusion discrepancy between marine and terrestrial protists highlights the difference in mechanisms involved in community assembly between these two environments. The novel network-phylogeny approach presented in this study have potential to unravel phylogenetic-driven assembly patterns in large scale datasets for which little is known about the taxonomy and function of the target organisms in other environments. The interplay between phylogeny and co-occurrence/co-exclusion networks remain to be disclosed in other microbial taxonomic groups, like Bacteria and Fungi, and among functional groups, like autotrophs, heterotrophs and associated microbes.

## Author contributions

GL and MD conceived the ideas; GL conducted the analyses; GL and MD wrote the manuscript.

## Acknowledgement

This work was supported by the Deutsche Forschungsgemeinschaft grants DU1319/5-1 to Micah Dunthorn. The authors are grateful to the High Performance Computer Elwetritsch at the Technical University of Kaiserslautern and the Centre for Computation of the Science Faculty of the University of Neuchâtel for computing support.

## Supporting information

**Figure S1.**
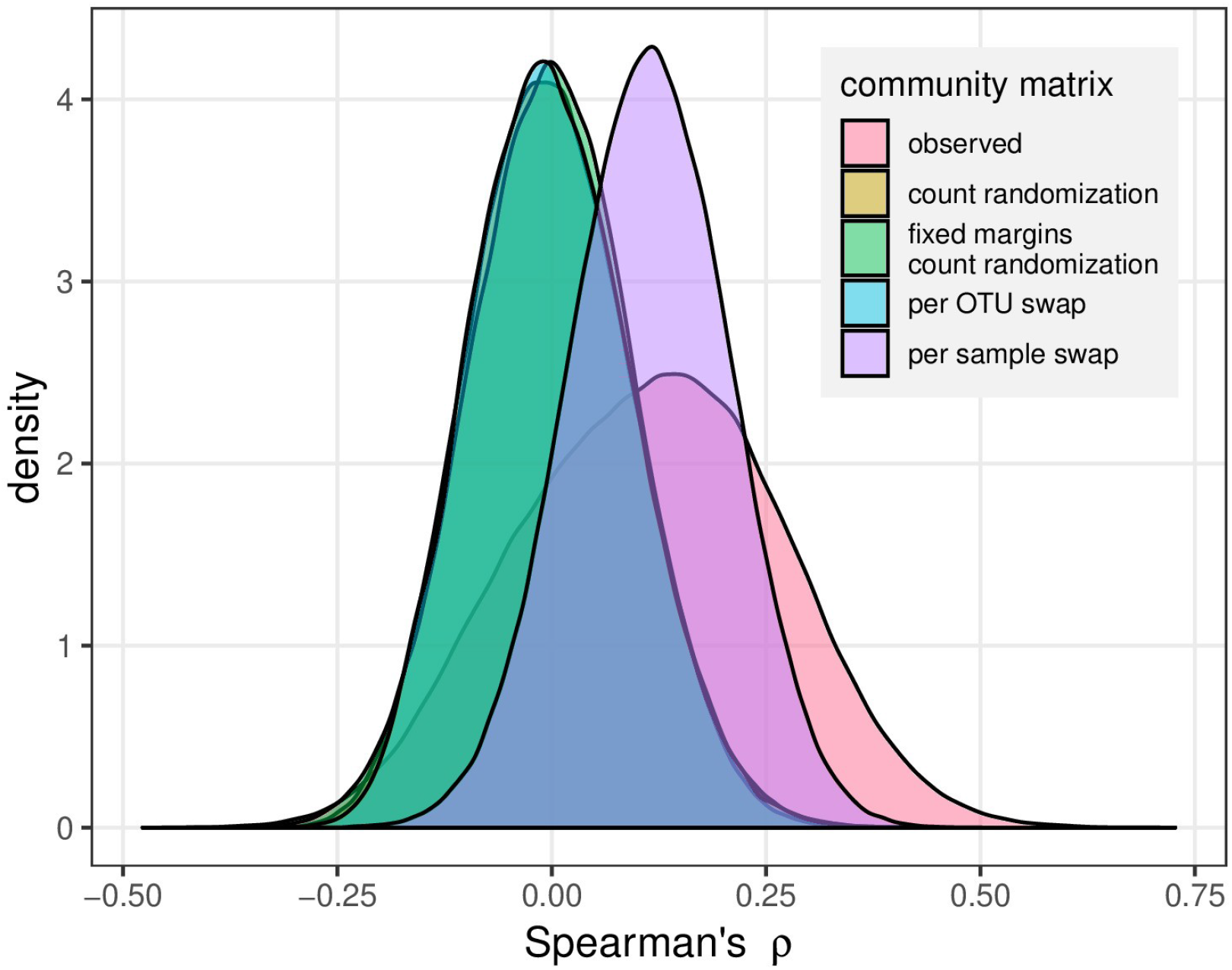
Distribution of Spearman’s *rho* correlations among OTUs normalized relative abundance calculated using the observed Neotropical soil community matrix (red) and four randomization of it: all counts were randomly drawn over the community matrix without constrain (yellow), all counts were randomly drawn while keeping OTU and samples sum fixed (green), abundance values were randomly swap within each OTU (blue), abundance values were randomly swap within each sample (purple). Overlapping yellow, green and blue areas produced a dark blue area with median and mean Spearman’s *rho* of zero. Observed and per sample swap matrices had median and mean Spearman’s *rho* of 0.11. The observed matrix had a standard deviation of 0.15, while this value was 0.09 for all randomized matrices.

**Figure S2.**
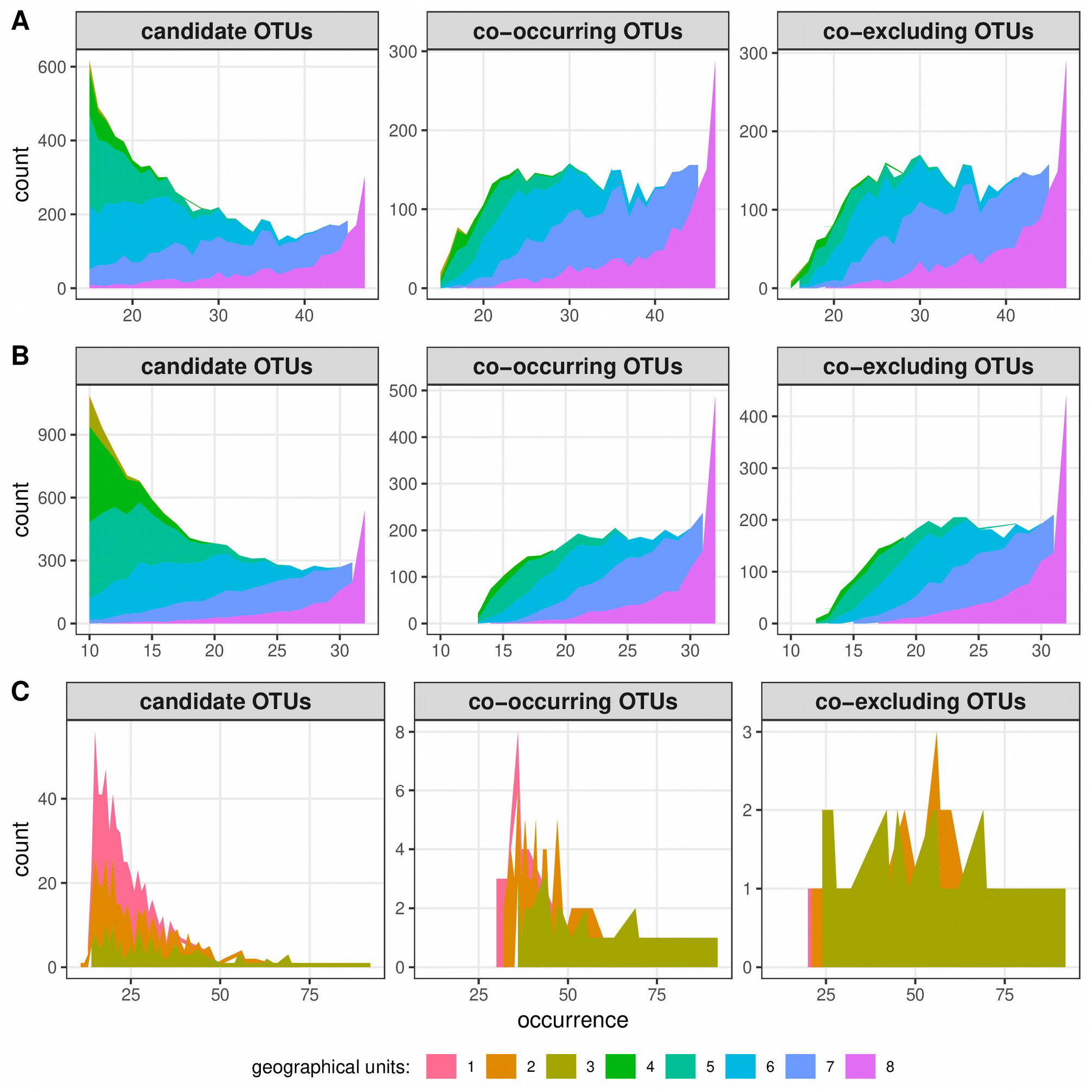
Distribution of candidate OTUs (OTUs occurring in at least 30 % of all samples for marine protists and in at least 10 % for terrestrial protists) and OTUs integrated into the cooccurrence or co-exclusion networks among the different geographical units: a. surface and b. DCM marine protist OTUs among eight different oceans and seas worldwide; c. Neotropical soil protist OTUs among three forests. Colored areas are for OTUs occurring in increasing amount of geographical units. Areas are stacked on each other (i.e. non-overlapping), so that the upper limit of the upper area is the cumulative amount of OTU occurring in the same number of samples.

**Figure S3.**
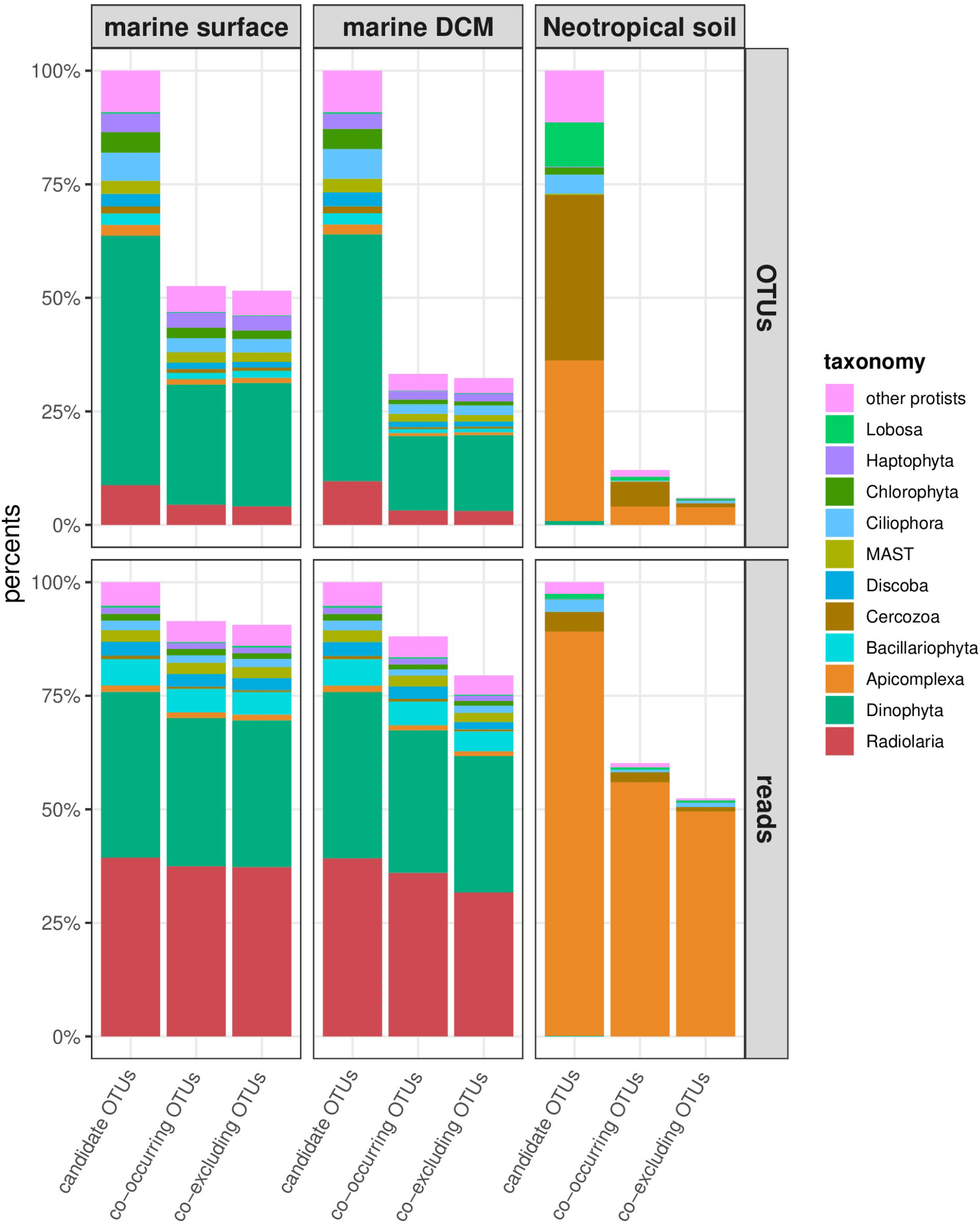
Taxonomy of candidate OTUs and OTUs integrated into the co-occurrence or coexclusion networks for each dataset, expressed in term of OTUs percentages and log-ratio transformed relative abundance percentages.

**Figure S4.**
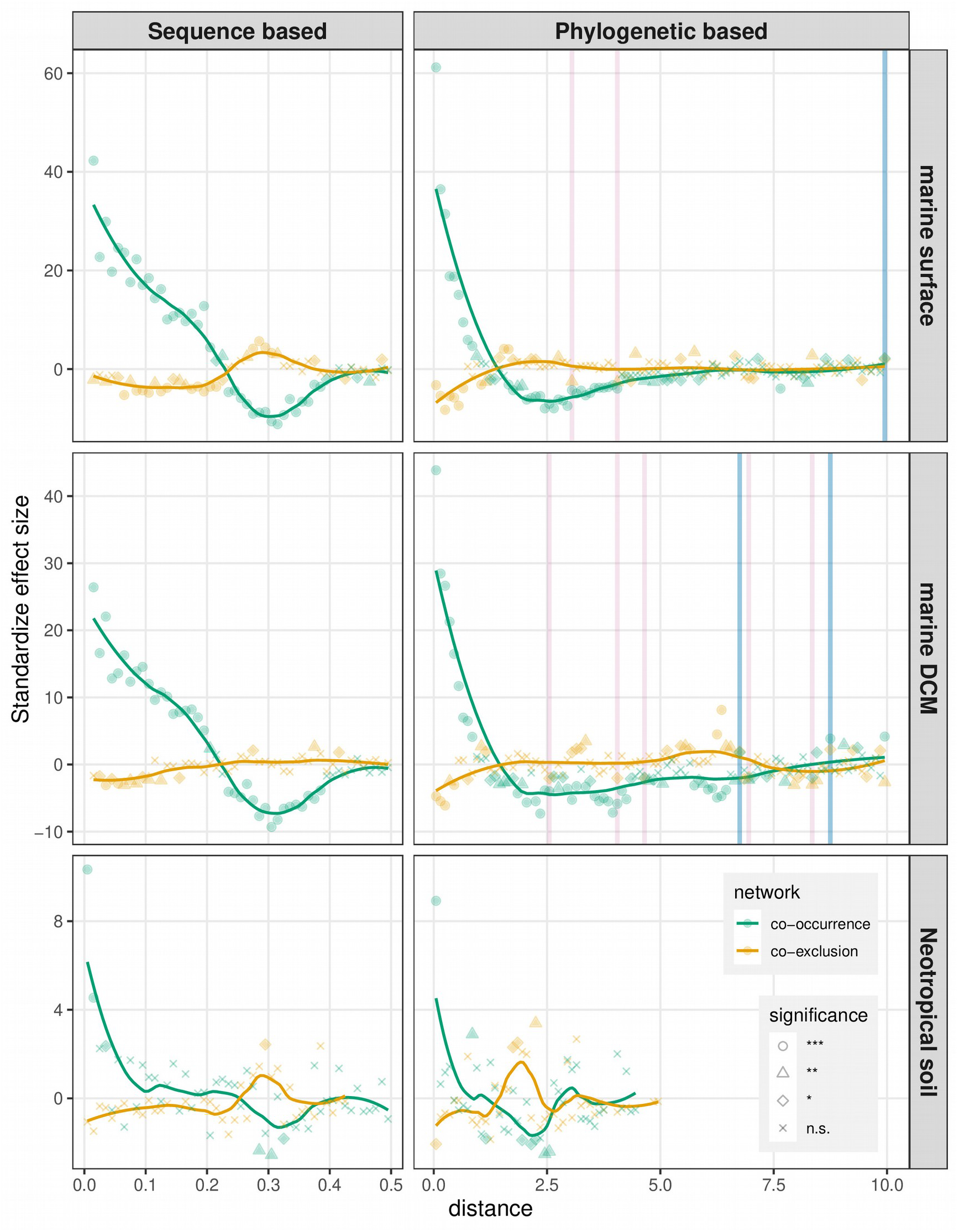
Standardize effect sizes (SES) in co-occurrence and co-exclusion networks compared to random networks with shuffled edges (null model 2). SES were calculated separately for stepwise increased pairwise sequence genetic distances and phylogenetic distances. The number of OTU pairs connected by an edge in the observed networks was accounted for each distance class (from 0 to 0.5 with a 0.01 step for sequence based; from 0 to the maximum phylogenetic distance with a 0.1 step for phylogenetic based) and compared to the corresponding distance class reported from the randomized networks. Two-sided non-parametric p.values are inversely proportional to the amount of null models with a higher (for positive SES) or lower (for negative SES) amount of cooccurrence than in the observed network for each distance class. P.values below or equal to 0.05 were considered significant (* ≤ 0.05; ** ≤ 0.01; *** ≤ 0.001). Distance ranges highlighted in blue or red are for distances with excess (significant positive SES) or lack (significant negative SES) of edges in both co-occurrence and co-exclusion networks simultaneously.

**Figure S5.**
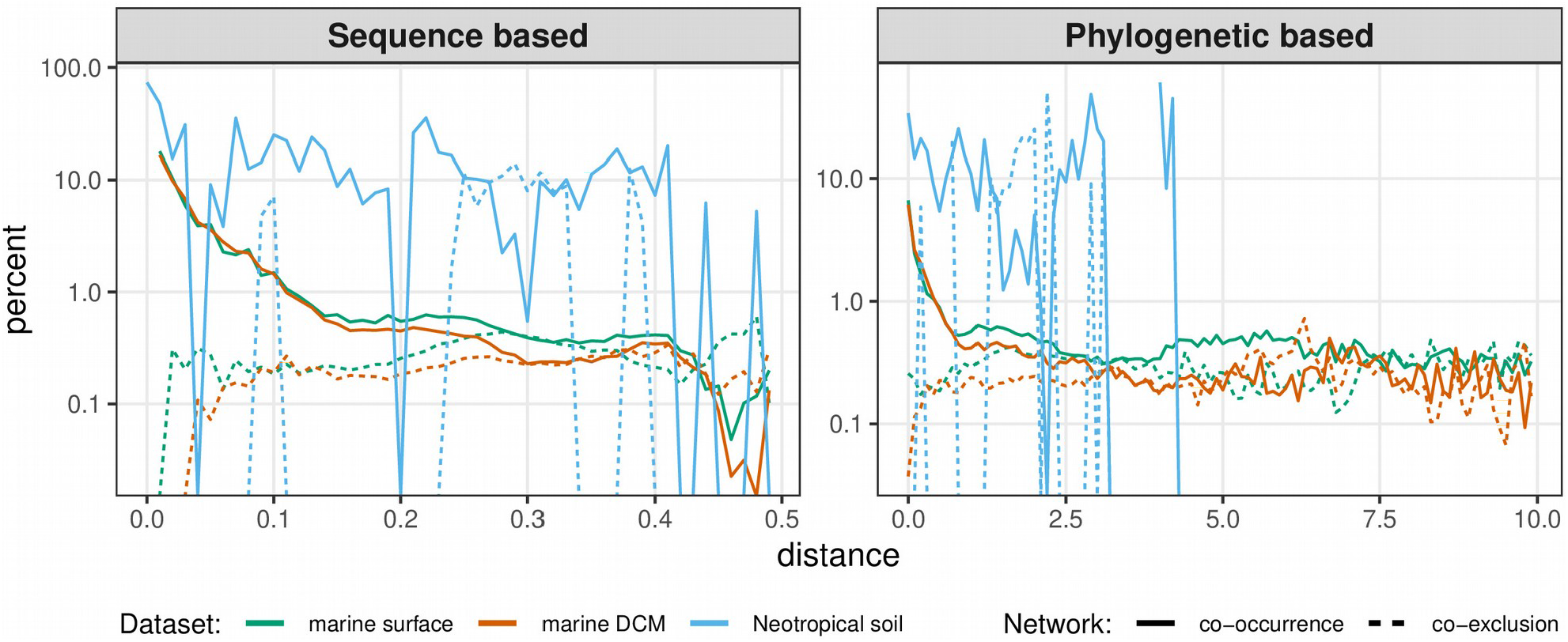
Percent of total candidate edges in the observed networks arranged by distance classes. Y-axis is square-root transformed to improve readability. The highest phylogenetic distance among candidate edges for Neotropical soil is 4.5. No slope were drawn for distance classes not covered in the observed networks.

**Figure S6.**
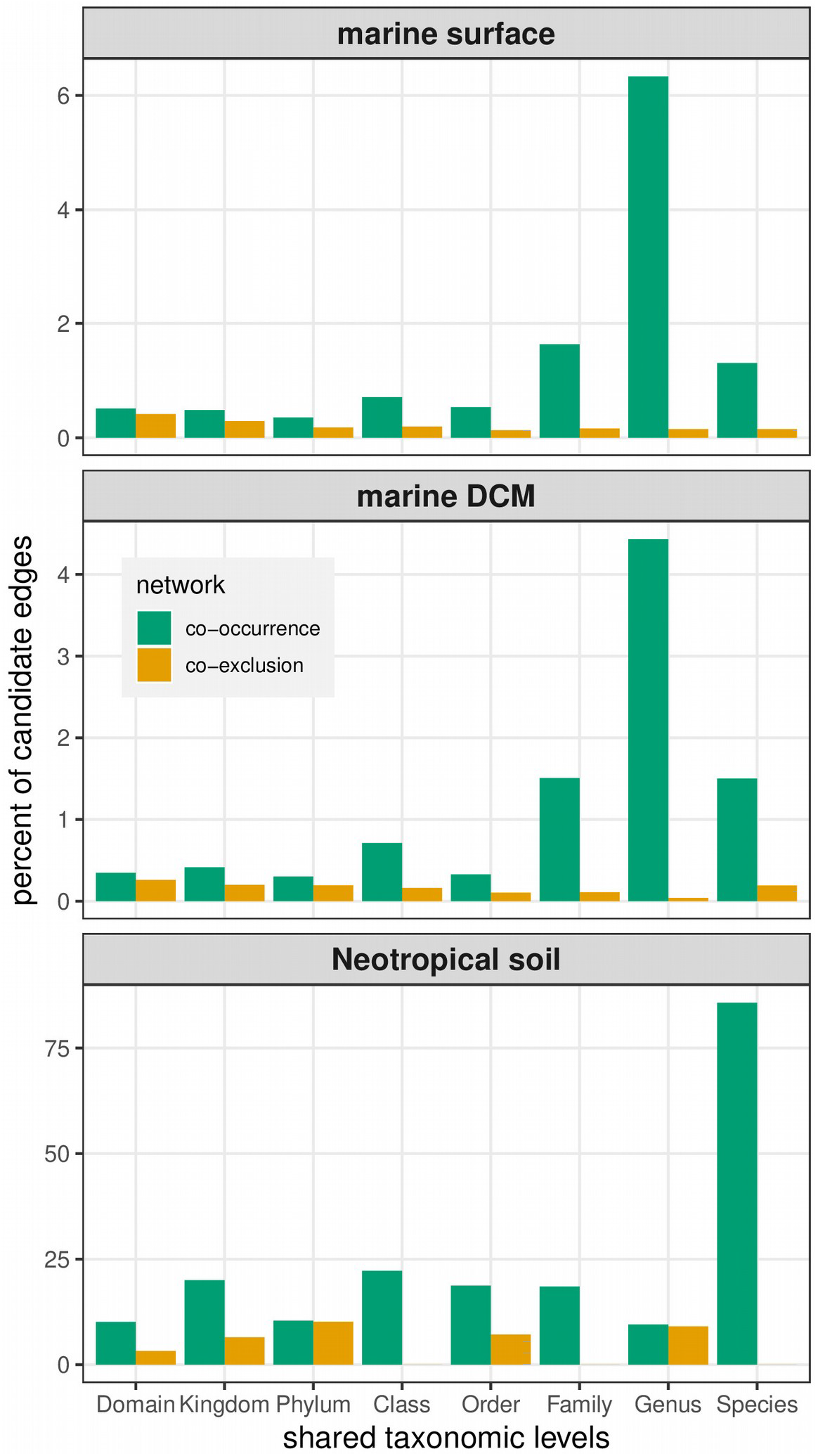
Percent of candidate edges sampled in the observed networks arranged by amount of shared taxonomic levels between co-occurring or co-excluding OTUs.

**Figure S7.**
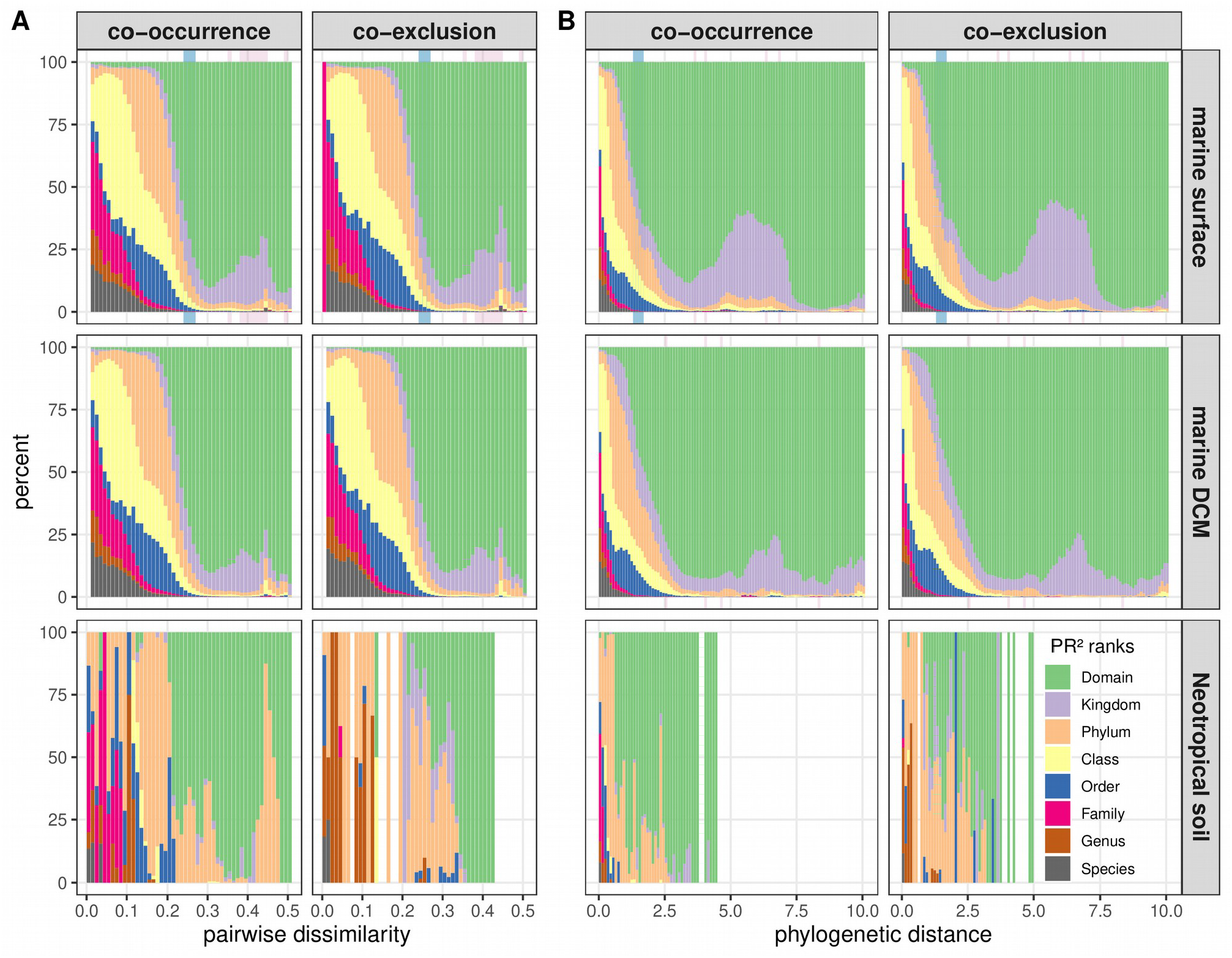
Distribution of taxonomic relationships between OTUs of all candidate edges for each pairwise sequence distance (a) and phylogenetic distance (b) classes. Blue and red shaded areas in the background are the distance classes with simultaneous positive or negative SES in both cooccurrence and co-exclusion networks using null model 1, as in Figure 2.

**Figure S8.**
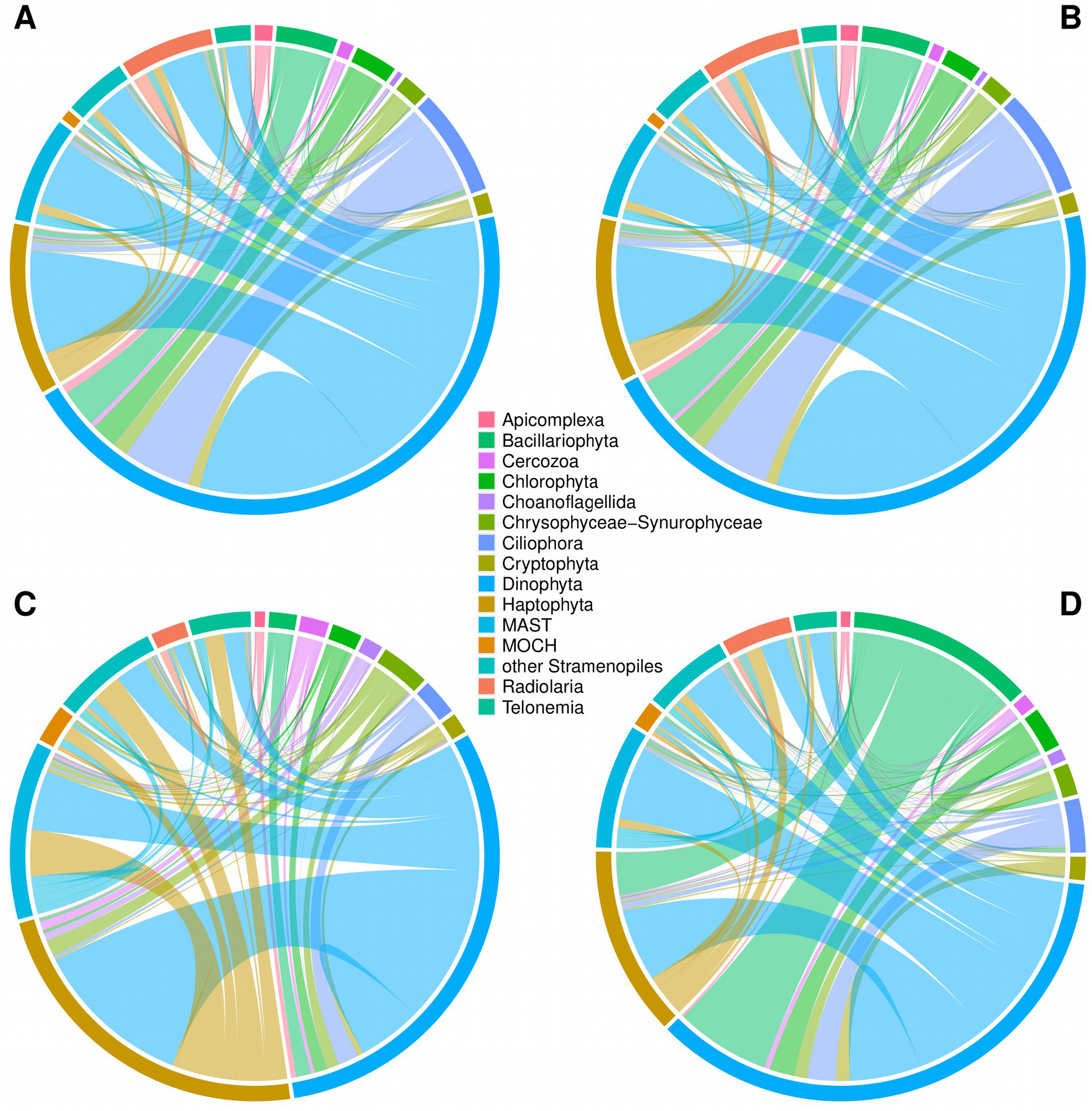
Proportion of edges between the different clades in the pairwise sequence genetic distance range 0.24-027 of the marine surface datasets. The two first chord diagrams represent all candidate edges for the co-occurrence (A) and co-exclusion (B) networks. The two last chord diagrams represent the observed distribution of edges in the co-occurrence (C) and co-exclusion (D) networks. Fold changes between observed and candidate edges ratio for each pair of clades are presented in Figure 4.

**Figure S9.**
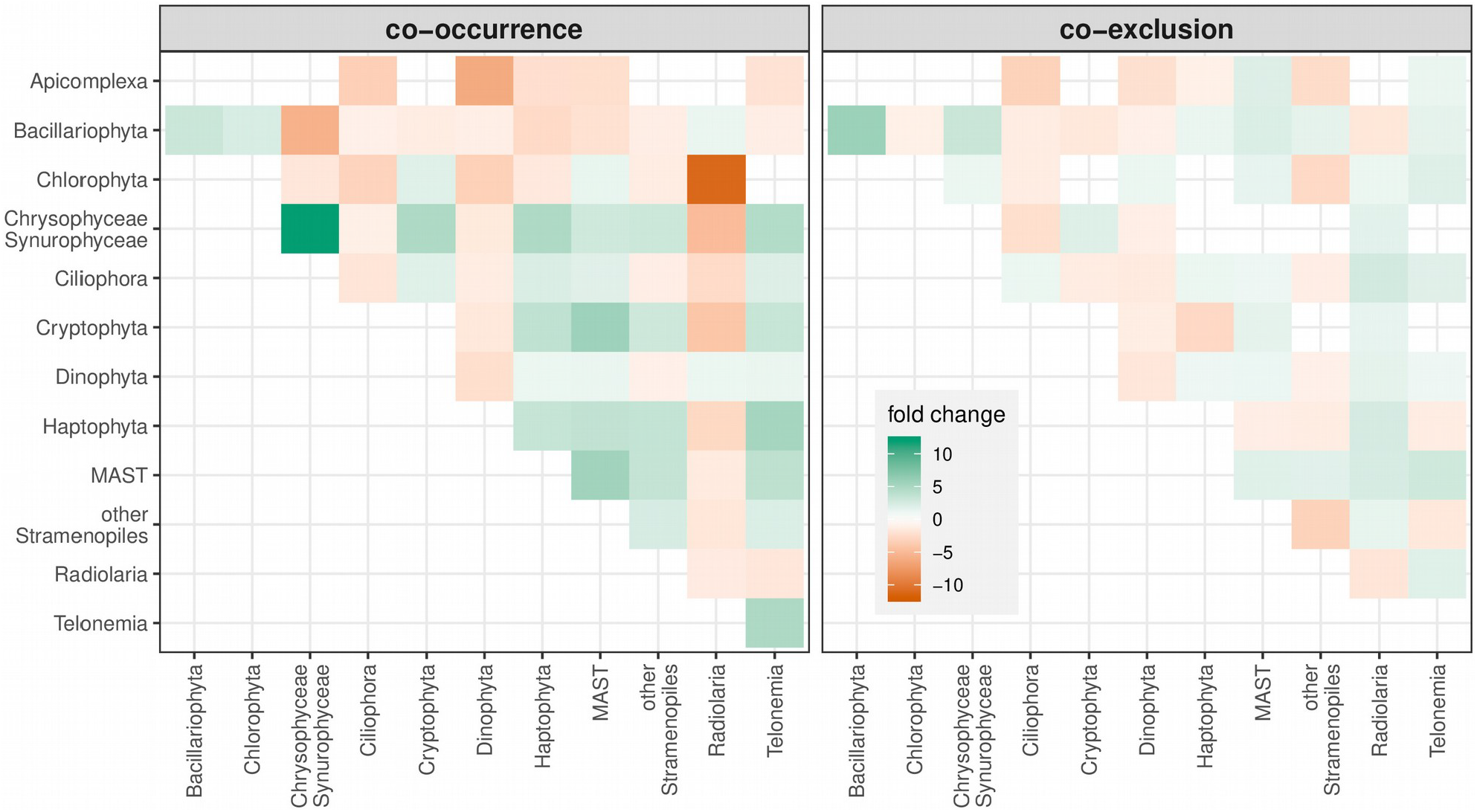
Fold changes in proportion of edges connecting the main clades in the marine DCM dataset compared to all candidate edges in the pairwise sequence distance range of 0.24-0.27 (*i.e*. the largest range of distance with simultaneous positive SES in co-occurrence and co-exclusion networks of the marine surface dataset when using the null model 1). The fold change color scale is identical to the one use for the marine surface dataset (Figure 4).

