## Supplementary material for "Relating network analyses to phylogenetic relatedness to infer protistan co-occurrences and co-exclusions in marine and terrestrial environments": File S1: File_S1.html

###### Guillaume Lentendu, Micah Dunthorn

- 1 Disclaimer
- 2 Prepare OTU tables
  - 2.1 Protist OTUs of the TARA Ocean dataset
  - 2.2 Protist OTUs of the Neotropical rainforest soils dataset
- 3 Networks
- 4 Phylogenies
- 5 Standardize effect sizes (SES.R)
- 6 submit to Slurm queue
- 7 Statistical analyses and figures
- 8 Dependencies
  - 8.1 bash
  - 8.2 R session informations


### 1 Disclaimer

This supplementary file provide the full statistical procedure used in this paper. Codes were executed on an HPC runing Debian 4.19.12-1 and Slurm queueing system and on a personal computer under Ubuntu 18.04.4. It might need to be adapted on other systems.

Bash code chunks begin with the shebang “#!/bin/bash” and are emphasized with blue borders, R codes are emphasized with orange borders.

### 2 Prepare OTU tables

#### 2.1 Protist OTUs of the TARA Ocean dataset

Source: de Vargas et al., 2015, http://doi.org/10.1126/science.1261605 as used in Mahé et al., 2017, http://doi.org/10.1038/s41559-017-0091

```
library(biomformat)
library(plyr)
library(tidyverse)
library(foreach)
library(iterators)

tara<-gzfile("Dataset_S1.biom.gz") %>% read_biom()
mat_raw<-t(as.matrix(biom_data(tara)))
ranks<-c("Domain","Kingdom","Phylum","Class","Order","Family","Genus","Species")
rank<-function(x) {ranks[as.numeric(sub(".*([0-9])$","\\1",x))]}
taxo_raw<-observation_metadata(tara) %>%
  rownames_to_column("OTU") %>%
  mutate_at(vars("total","identity"),list(as.numeric)) %>%
  mutate_at(vars(starts_with("taxonomy")),list(as.factor)) %>%
  rename_at(vars(starts_with("taxonomy")),list(rank))

env_raw<-sample_metadata(tara) %>%
  rownames_to_column("sample") %>%
  mutate_at(vars("depth","biome","region"),list(as.factor))

# keep only Surface and DCM samples
env_ok<-filter(env_raw,depth %in% c("Surface","Deep_Chlorophyl_Maximum")) %>% droplevels()
mat_ok<-mat_raw[env_ok$sample,] %>% .[,which(colSums(.)>0)]
taxo_ok<-filter(taxo_raw,OTU %in% colnames(mat_ok))
rm(tara,mat_raw,env_raw,taxo_raw)

# Pool libraries of identical station and depth (i.e. pool all size fractions, always one library type only per size fraction)
env_merged<-group_by(env_ok,station,depth,biome,region) %>%
  summarize(samples=paste(sample,collapse="_")) %>%
  mutate(abb=mapvalues(depth,levels(depth),c("DCM","SUR")),
         name=paste(station,abb,sep="_"))
mat_merged<-foreach(i=iapply(env_merged,1),.combine=rbind) %do% {
  if(grepl("_",i$samples)) {
    t(setNames(data.frame(colSums(mat_ok[unlist(strsplit(as.character(i$samples),"_")),])),i$name))
  } else {
    t(setNames(data.frame(mat_ok[as.character(i$samples),]),i$name))
  }
}

# removed unidentified eukaryotes/Opisthokonta OTUs (potential macroorganisms)
taxo<-filter(taxo_ok,Kingdom!="Eukaryota_X",Phylum!="Opisthokonta_X")
mat<-mat_merged[,taxo$OTU]

# Split between the 2 depths and export to tsv or RDS
for (i in levels(env_merged$abb)) {
  tmp_env<-filter(env_merged,abb==i)
  assign(paste0("env_",i),tmp_env)
  saveRDS(tmp_env,paste0("tara_",i,"_env"))
  tmp_mat<-mat[tmp_env$name,] %>% .[,which(colSums(.)>0)]
  assign(paste0("mat_",i),tmp_mat)
  write.table(tmp_mat,paste0("tara_",i,"_mat.tsv"),sep="\t",quote=F)
  tmp_taxo<-filter(taxo,OTU %in% colnames(tmp_mat)) %>%
    droplevels()
  assign(paste0("taxo_",i),tmp_taxo)
  saveRDS(tmp_taxo,paste0("tara_",i,"_taxo"))
}
```

#### 2.2 Protist OTUs of the Neotropical rainforest soils dataset

Source: Mahé et al., 2017, http://doi.org/10.1038/s41559-017-0091

```
library(biomformat)
library(plyr)
library(tidyverse)
library(foreach)
library(iterators)

neotrop<-gzfile("Dataset_S2.biom.gz") %>% read_biom()
mat<-t(as.matrix(biom_data(neotrop)))
ranks<-c("Domain","Kingdom","Phylum","Class","Order","Family","Genus","Species","Genus2","Species2")
rank<-function(x) {ranks[as.numeric(sub("^[a-z]*([0-9][0-9]*)$","\\1",x))]}
taxo<-observation_metadata(neotrop) %>%
  rownames_to_column("OTU") %>%
  mutate_at(vars("total","identity"),list(as.numeric)) %>%
  mutate_at(vars(starts_with("taxonomy")),~sub("\\*","unclassified",.)) %>% 
  mutate_at(vars(starts_with("taxonomy")),list(as.factor)) %>%
  rename_at(vars(starts_with("taxonomy")),list(rank))

env<-sample_metadata(neotrop) %>%
  rownames_to_column("sample") %>%
  mutate_at(vars("ab","station","forest","date"),list(as.factor)) %>%
  mutate_at(vars("longitude","latitude"),list(as.numeric))

# merge single and pooled samples of the same location (i.e. sum reads)
to_merge<-NULL
for(i in 1:nrow(mat)) {
  for(j in 1:nrow(mat)) {
    if(i<j) {
      if(grepl(rownames(mat)[j],rownames(mat)[i])){
        to_merge<-rbind(to_merge,c(rownames(mat)[i],rownames(mat)[j]))
      }}}}
my_merge<-function(x) {
  full_join(summarize(x,sample=paste(sample,collapse=",")),
            summarize_at(x,vars(-sample),list(first)))
}
env_merged<-full_join(env,setNames(data.frame(to_merge),c("name","sample"))) %>%
  mutate(name=ifelse(is.na(name),sample,as.character(name))) %>%
  group_by(name) %>%
  my_merge(.)
mat_merged<-foreach(i=iapply(env_merged,1),.combine=rbind) %do% {
  if(grepl(",",i$sample)) {
    t(setNames(data.frame(colSums(mat[unlist(strsplit(i$sample,",")),])),i$name))
  } else {
    t(setNames(data.frame(mat[i$sample,]),i$name))
  }
}

# removed unidentified eukaryotes/Opisthokonta OTUs (potential macroorganisms)
taxo_ok<-filter(taxo,! Phylum %in% c("unclassified","Opisthokonta_X")) %>%
  droplevels()
mat_ok<-mat_merged[,taxo_ok$OTU]

# export to tsv
write.table(mat_ok,"neotropical_mat.tsv",sep="\t",quote=F)
saveRDS(taxo_ok,"neotropical_taxo")
saveRDS(env_merged,"neotropical_env")
```

### 3 Networks

Following Connor, Barbéran and Clasuet (2017, http://doi.org/10.1371/journal.pone.0176751), as re-implemented in the tool NetworkNullHPC (https://github.com/lentendu/NetworkNullHPC)

```
#!/bin/bash

NetworkNullHPC.sh -m 1 -n ratio_log -o 0.3 tara_SUR_mat.tsv
NetworkNullHPC.sh -m 1 -n ratio_log -o 0.3 tara_DCM_mat.tsv
NetworkNullHPC.sh -m 1 -n ratio_log -r 0.2 neotropical_mat.tsv
```

### 4 Phylogenies

Retrieve nucleotide sequences from OTUs included in the networks

network\_fasta.R :

```
library(tidyverse)

project<-commandArgs()[7]
taxo<-readRDS(paste0(project,"_taxo"))

# candidate OTUs representative sequences
nnhpc<-paste(unlist(strsplit(list.files(patter=paste0("^info.*",project,"_mat.txt")),"\\."))[2:3],collapse=".")
candidate_otus<-scan(file.path(nnhpc,"otus"),what='character')
candidate_taxo<-filter(taxo,OTU %in% candidate_otus) %>%
  select(OTU,sequence)

# network OTUs representative sequences
cooc<-read.table(file.path(paste0("cooccurrence.",strsplit(nnhpc,"\\.")[[1]][2],".",project,"_mat.txt")),
                      col.names=c("from","to","cor"),stringsAsFactors=F)
cooc_otus<-unique(c(cooc$from,cooc$to))
coex<-read.table(file.path(paste0("coexclusion.",strsplit(nnhpc,"\\.")[[1]][2],".",project,"_mat.txt")),
                 col.names=c("from","to","cor"),stringsAsFactors=F)
coex_otus<-unique(c(coex$from,coex$to))

network_taxo<-filter(candidate_taxo,OTU %in% cooc_otus | OTU %in% coex_otus) %>%
  droplevels()

write(sub("^",">",apply(network_taxo,1,paste,collapse="\n")),
      paste0(project,".fasta"),ncolumns=1)
```

Prepare submission script for pairwise alignments, multiple alignments and ML tree inferences

```
#!/bin/bash

dataset=(neotropical tara_SUR tara_DCM)
mkdir phylogeny logfiles

# utility script
cat <<EOF > fas2phylip
#!/bin/bash
sed '/>/G' \$@ |  sed -e :a -e '\$!N;/>/!s/\n//;ta' -e 'P;D' | sed 's/^>//;\$!N;s/\n/\t/' | awk 'NR==FNR{len=length(\$2); a[NR]=\$0}END{print FNR,len; for(i=1;i<=FNR;i++){print a[i]}}'
EOF
chmod +x fas2phylip

# loop to prepare submission scripts
for i in ${dataset[@]}
do

# pairwise distances
    cat << EOF > sub_pairwise_$i
#!/bin/bash

#SBATCH -J pairwise_${i}
#SBATCH -o logfiles/pairwise_${i}.%j.out
#SBATCH -e logfiles/pairwise_${i}.%j.err
#SBATCH -t 12:00:00
#SBATCH -N 1
#SBATCH -n 1
#SBATCH --cpus-per-task=16
#SBATCH --mem=32G

module load R/3.5.1 sumatra/1.0.31

# fasta file of network OTUs
Rscript --vanilla network_fasta.R ${i}

# pairwise distances
sumatra -d -p \$SLURM_CPUS_PER_TASK ${i}.fasta | awk '{gsub("OTU_","");if(\$2<\$1){print "OTU_"\$2,"OTU_"\$1,\$3} else print "OTU_"\$1,"OTU_"\$2,\$3}' > ${i}.suma.pairwise
EOF

# MAFFT
        cat << EOF > sub_mafft_${i}
#!/bin/bash

#SBATCH -J mafft_${i}
#SBATCH -o logfiles/mafft_${i}.%j.out
#SBATCH -e logfiles/mafft_${i}.%j.err
#SBATCH -t 48:00:00
#SBATCH -N 1
#SBATCH -n 1
#SBATCH --cpus-per-task=16
#SBATCH --mem=8G

cd phylogeny
mkdir ${i}
module load mafft/7.407
mafft --retree 2 --maxiterate 2 --thread \$SLURM_CPUS_PER_TASK ../${i}.fasta > ${i}.FFT-NS-i.mafft
../fas2phylip ${i}.FFT-NS-i.mafft > ${i}/${i}.FFT-NS-i.mafft.phy
EOF

# RAxML
        cat << EOF > sub_raxml_${i}
#!/bin/bash

#SBATCH -J raxml_${i}
#SBATCH -o logfiles/raxml_${i}.%j.out
#SBATCH -e logfiles/raxml_${i}.%j.err
#SBATCH -L ompi
#SBATCH -N 16
#SBATCH -n 256
#SBATCH --ntasks-per-node=16
#SBATCH -t 48:00:00
#SBATCH --mem=32G

cd phylogeny/${i}
module load raxml/8.2.11
mpirun raxmlHPC-MPI-AVX -f d -m GTRCAT -s ${i}.FFT-NS-i.mafft.phy -p 12345 -# \$SLURM_NTASKS -n MultipleOriginal_${i}
EOF

done
```

### 5 Standardize effect sizes (SES.R)

```
library(igraph)
library(doParallel)
library(ape)
library(picante)
library(plyr)
library(tidyverse)
library(ggplot2)
library(abind)
library(scales)

# read input
project<-commandArgs()[7]
ncores<-as.numeric(commandArgs()[8])
type<-commandArgs()[9]
ty<-substr(type,1,4)

# load networks
net<-read.table(list.files(pattern=paste0(type,".[0-9]*.",project,"_mat.txt")),
                 stringsAsFactors=F,col.names=c("from","to","cor")) %>%
  graph_from_data_frame(directed=F)
net_name<-data.frame(get.edgelist(net),stringsAsFactors=F) %>%
  setNames(c("from","to")) %>%
  unite(pair,c("from","to"),sep="-")
net_node_name<-V(net)$name
net_degree<-degree(net)
net_nbedges<-length(E(net))

# load phylogenetic tree
tree<-read.tree(file.path("phylogeny",project,paste0("RAxML_bestTree.MultipleOriginal_",project)))
net_tree<-drop.tip(tree,data.frame(tip=tree$tip.label,stringsAsFactors=F) %>%
                      filter(! tip %in% V(net)$name) %>% .$tip)

# get all pairwise sequence disimilarities and calculate all phylogenetic distances
pair_dist<-read.table(paste0(project,".suma.pairwise"),col.names=c("from","to","dist"),stringsAsFactors=F) %>%
  mutate(pair=paste(from,to,sep="-")) %>%
  filter(from %in% net_node_name,to %in% net_node_name) %>%
  select(pair,dist)

phylo_dist<-setNames(data.frame(t(combn(net_tree$tip.label,2)),c(as.dist(cophenetic(net_tree))),
                                stringsAsFactors=F),
                     c("from","to","dist")) %>%
  mutate(pair=ifelse(from<to,paste(from,to,sep="-"),paste(to,from,sep="-"))) %>%
  select(pair,dist)

# observe network distances (i.e. only for edges in the network)
obs_dist<-rbind(data.frame(method="Sequence dissimilarity",digits=2,max=0.49,pair_dist),
                data.frame(method="Phylogenetic distance",digits=1,max=9.9,phylo_dist)) %>%
  right_join(net_name)
obs_dist_attr<-select(obs_dist,method,dist,pair) %>%
  mutate(method=mapvalues(method,levels(method),c("pairwise","phylo"))) %>%
  spread(method,dist)
for (i in c("pairwise","phylo")) {
  edge_attr(net,i)<-obs_dist_attr[[i]]
}
saveRDS(net,paste(project,ty,"network",sep="_"))

# Stats
obs_dist_stat<-dlply(obs_dist,.(method),function(x) {
  mutate(x,dist=factor(ifelse(dist>max,max,floor(dist*10^digits)*10^-digits),
                       levels=seq(0,unique(max),10^(-unique(digits))))) %>%
    group_by(dist) %>%
    summarize(obs=n()) %>%
    complete(dist,fill=list(obs=0))
})

# proportion of all potential network distances covered in the observed network
full_dist<-rbind(data.frame(method="Sequence dissimilarity",digits=2,max=0.49,pair_dist),
                 data.frame(method="Phylogenetic distance",digits=1,max=9.9,phylo_dist)) %>%
  ddply(.(method),function(x) {
    mutate(x,dist=factor(ifelse(dist>max,max,floor(dist*10^digits)*10^-digits),
                         levels=seq(0,unique(max),10^(-unique(digits))))) %>%
      group_by(dist) %>%
      summarize(full=n()) %>%
      complete(dist,fill=list(full=0))
  }) %>%
  mutate(full=ifelse(full==0,NA,full)) %>%
  left_join(ldply(obs_dist_stat)) %>%
  mutate(percent=obs/full*100)
saveRDS(full_dist,paste(project,ty,"network_coverage",sep="_"))

# Network and phylogenetic null models:
# network based: shuffle edges, output edge names, to left_join with obs_dist
# tree based: shuffle tip of tree, output in-order pair names, sliced with the position of network edges found in obs_dist, to left_join with obs_dist
mapcbind<-function(...){Map(cbind,...)}
cl<-makeCluster(ncores)
registerDoParallel(cl)
rand_pairs<-foreach(x=1:ncores,.combine=mapcbind,.multicombine=T,.inorder=T,.packages=c('foreach','plyr','dplyr','tidyr','igraph','picante')) %dopar% {
  my_seq<-floor(seq(1,1001,length.out=ncores+1))
  foreach(y=my_seq[x]:(my_seq[x+1]-1),.combine=mapcbind,.multicombine=T) %do% {
    set.seed(y)
    tmp_net<-sample_fitness(net_nbedges,net_degree)
    V(tmp_net)$name<-net_node_name
    set.seed(y)
    tmp_tree<-tipShuffle(net_tree)
    tmp_name<-paste0("null_",y)
    tmp_pair_net<-setNames(data.frame(get.edgelist(tmp_net),stringsAsFactors=F),nm=c("from","to")) %>%
      mutate(f=as.numeric(sub("OTU_","",from)),t=as.numeric(sub("OTU_","",to)),
             pair=ifelse(f<t,paste(from,to,sep="-"),paste(to,from,sep="-")))
    tmp_net_pair_dist<-left_join(tmp_pair_net,pair_dist)$dist
    tmp_net_phylo_dist<-left_join(tmp_pair_net,phylo_dist)$dist
    tmp_pair_tree<-setNames(data.frame(t(combn(tmp_tree$tip.label,2))),c("from","to")) %>%
      mutate(f=as.numeric(sub("OTU_","",from)),t=as.numeric(sub("OTU_","",to)),
             pair=ifelse(f<t,paste(from,to,sep="-"),paste(to,from,sep="-")))
    tmp_tree_pair_dist<-pair_dist[match(net_name$pair,tmp_pair_tree$pair),"dist"]
    tmp_tree_phylo_dist<-phylo_dist[match(net_name$pair,tmp_pair_tree$pair),"dist"]
    tmp_df<-rbind(cbind.data.frame(method="Sequence dissimilarity",digits=2,max=0.49,
                           rbind(cbind.data.frame(model="random network",dist=tmp_net_pair_dist),
                                 cbind.data.frame(model="shuffled tree tips",dist=tmp_tree_pair_dist))),
          cbind.data.frame(method="Phylogenetic distance",digits=1,max=9.9,
                           rbind(cbind.data.frame(model="random network",dist=tmp_net_phylo_dist),
                                 cbind.data.frame(model="shuffled tree tips",dist=tmp_tree_phylo_dist)))) 
    tmp_df_stat<-dlply(tmp_df,.(method,model),function(x) {
      tmp_x<-mutate(x,dist=factor(ifelse(dist>max,max,floor(dist*10^digits)*10^-digits),levels=seq(0,unique(max),10^(-unique(digits))))) %>%
        group_by(dist) %>% summarize(null=n()) %>% complete(dist,fill=list(null=0)) %>% rename(!!tmp_name:=null)
      if(y==1) {return(select(tmp_x,dist,starts_with("null")))}
      else {return(select(tmp_x,starts_with("null")))}
      })
  }
}
stopCluster(cl)

# merge obs and null data together
all_dist_stat<-right_join(ldply(obs_dist_stat),
                          ldply(rand_pairs,.id="name") %>% separate(name,c("method","model"),sep="\\.") %>%
                            mutate(method=factor(method,levels=unique(method)),model=as.factor(model)))

# standardize effect size
ses<-cbind.data.frame(all_dist_stat,mean=rowMeans(select(all_dist_stat,starts_with("null"))),
                      sd=apply(select(all_dist_stat,starts_with("null")),1,sd),
                      sup=apply(select(all_dist_stat,obs,starts_with("null")),1,function(x) length(which(x[2:length(x)]>x[1]))),
                      inf=apply(select(all_dist_stat,obs,starts_with("null")),1,function(x) length(which(x[2:length(x)]<x[1])))) %>%
  mutate(ses=(obs-mean)/sd,pval=ifelse(obs<mean,1-sup/1000,1-inf/1000),
         signif=factor(pval<=0.05,levels=c("TRUE","FALSE"))) %>%
  select( -starts_with("null")) %>%
  group_by(method) %>%
  arrange(method) %>%
  mutate(dist=as.numeric(as.character(dist)),
         dist=dist+5*10^-(1+nchar(sub("^.*\\.","",max(dist)))),
         size=0.8+0.8*(100-length(unique(dist)))/50)
saveRDS(ses,paste(project,ty,"ses_df",sep="_"))
```

### 6 submit to Slurm queue

```
#!/bin/bash

dataset=(neotropical tara_SUR tara_DCM)

# SES submission scripts
for i in ${dataset[@]}
do
    cat << EOF > sub_SES_$i
#!/bin/bash

#SBATCH -J ${i}_SES
#SBATCH -o logfiles/${i}_SES.%j.out
#SBATCH -e logfiles/${i}_SES.%j.err
#SBATCH -t 48:00:00
#SBATCH -N 1
#SBATCH -n 1
#SBATCH --cpus-per-task=16
#SBATCH --mem=256G

module load R/3.5.1
Rscript --vanilla SES.R ${i} \$SLURM_CPUS_PER_TASK cooccurrence
Rscript --vanilla SES.R ${i} \$SLURM_CPUS_PER_TASK coexclusion
EOF
done

# Submit to queue

declare -A jobid_pairwise jobid_mafft jobid_raxml jobid_SES
for i in ${dataset[@]}
do
    jobid_pairwise[$i]=$(sbatch --parsable sub_pairwise_${i})
    jobid_mafft[$i]=$(sbatch --parsable -d afterok:${jobid_pairwise[$i]} sub_mafft_${i})
    jobid_raxml[$i]=$(sbatch --parsable -d afterok:${jobid_mafft[$i]} sub_raxml_${i})
    jobid_SES[$i]=$(sbatch --parsable -d afterok:${jobid_raxml[$i]} sub_SES_$i)
    echo "job submitted for dataset ${i}:"
    echo "${jobid_pairwise[$i]} ${jobid_mafft[$i]} ${jobid_raxml[$i]} ${jobid_SES[$i]}"
done
```

### 7 Statistical analyses and figures

```
library(vegan)
library(Hmisc)
library(ape)
library(circlize)
library(ComplexHeatmap)
library(igraph)
library(ggpubr)
library(ggplot2)
library(ggnetwork)
library(scales)
library(ggthemes)
library(grid)
library(plyr)
library(tidyverse)
library(foreach)
library(iterators)

# set plot theme
theme_set(theme_bw())
theme_update(panel.grid.minor=element_blank(),
             strip.text=element_text(size=12,face="bold"),
             axis.title=element_text(size=12),
             legend.background=element_rect(fill="grey95"),
             legend.text=element_text(size=8),
             legend.title=element_text(size=10),
             legend.key.size=unit(1,"line"),
             legend.box.just="right")

# list files
files<-sub("_ses_df","",list.files("./",pattern="*_ses_df")) %>%
  data.frame(file=.,stringsAsFactors=F) %>%
  mutate(project=as.factor(sub("_co[a-z]*","",file)),
         type=as.factor(sub(".*_(co[a-z]*)","\\1",file))) %>%
  left_join(data.frame(net=sub("_mat\\.txt","",list.files(pattern="co[a-z]*\\.[0-9]*\\..*_mat.txt"))) %>%
              separate(net,c("type","nnhpc","project"),sep="\\.") %>%
              mutate(type=as.factor(substr(type,1,4)),
                     nnhpc=as.numeric(nnhpc),
                     project=as.factor(project)))

# load networks, OTU matrices and taxonomy tables
for(i in files$file) {
  assign(paste0(i,"_net"),readRDS(paste0(i,"_network")))
}
for (i in levels(files$project)) {
  tmp_mat<-readRDS(file.path(paste0("NetworkNullHPC.",filter(files,project==i,type=="cooc")$nnhpc),"mat"))
  rownames(tmp_mat)<-sub("_[A-Z]*$","",rownames(tmp_mat))
  assign(paste0(i,"_mat"),tmp_mat)
  tmp_taxo<-readRDS(paste0(i,"_taxo")) %>%
    filter(OTU %in% colnames(tmp_mat))
  assign(paste0(i,"_taxo"),tmp_taxo)
  tmp_env<-readRDS(paste0(i,"_env")) %>%
    ungroup()
  if (i=="neotropical") {
    tmp_env<-select(tmp_env,name,forest) %>%
      rename(sample=name,unit=forest) %>%
      filter(sample %in% rownames(tmp_mat))
  } else {
    tmp_env<-select(tmp_env,station,region) %>%
      rename(sample=station,unit=region)
  }
  assign(paste0(i,"_env"),tmp_env)
}

##########################################
## example Spearman's rho distributions ##
##########################################
# for the Neotropical soil dataset
seed<-12345
# neotropical_mat = normalized matrix
obs<-neotropical_mat
# count randomization without fixed margins
set.seed(seed); rand0<-permatfull(neotropical_mat,fixedmar="none",times=1)$perm[[1]]
# count randomization with fixed margins (computational intensive)
set.seed(seed); rand1<-permatfull(neotropical_mat,fixedmar="both",times=1)$perm[[1]]
# swap with fixed OTU margin randomization
set.seed(seed); rand2<-apply(neotropical_mat,2,sample)
# swap with fixed sample margin randomization
set.seed(seed); rand3<-t(apply(neotropical_mat,1,sample))

b<-1e-4
spear<-ldply(as.list(setNames(c("obs",paste0("rand",0:3)),c("obs",paste0("rand",0:3)))),function(i){
  # noise addition
  set.seed(seed); noise<-get(i)+(-b+(2*b)*matrix(runif(length(c(get(i)))),nrow=nrow(get(i))))
  # re-normalize per sample if necessary
  if(sd(rowSums(noise))/ncol(noise)>1) {
    noise_norm<-round(noise*(median(rowSums(noise))*0.5/rowSums(noise)))
  } else {
    noise_norm<-noise
  }
  #spearman's rho
  data.frame(cor=c(as.dist(rcorr(noise_norm,type="spearman")$r)))
},.id="matrix") %>%
  mutate(matrix=as.factor(matrix),
         matrix=mapvalues(matrix,levels(matrix),c("observed","count randomization","fixed margins\ncount randomization","per OTU swap","per sample swap")))

plot_cor<-ggplot(spear,aes(cor)) +
  geom_density(aes(fill=matrix),alpha=0.5) +
  theme(legend.position=c(0.8,0.75)) +
  labs(x=expression("Spearman's "~italic(rho)),fill="community matrix") +
  scale_fill_hue(h=c(0,280))
ggsave("Figure_S1.pdf",plot_cor,width=5,height=4)

#######################################
## example networks with null models ##
#######################################
# simplify example network: only the largest component
net<-induced_subgraph(neotropical_cooc_net,which(components(neotropical_cooc_net)$membership==3))

# load phylogenetic tree
tree<-read.tree("phylogeny/neotropical/RAxML_bestTree.MultipleOriginal_neotropical")
net_tree<-drop.tip(tree,data.frame(tip=tree$tip.label,stringsAsFactors=F) %>%
                     filter(! tip %in% V(net)$name) %>% .$tip)

# get all pairwise sequence disimilarities and calculate all phylogenetic distances
pair_dist<-read.table("neotropical.suma.pairwise",col.names=c("from","to","dist"),stringsAsFactors=F) %>%
  mutate(pair=paste(from,to,sep="-")) %>%
  filter(from %in% V(net)$name,to %in% V(net)$name) %>%
  select(pair,dist)
phylo_dist<-setNames(data.frame(t(combn(net_tree$tip.label,2)),c(as.dist(cophenetic(net_tree))),
                                stringsAsFactors=F),
                     c("from","to","dist")) %>%
  mutate(pair=ifelse(from<to,paste(from,to,sep="-"),paste(to,from,sep="-"))) %>%
  select(pair,dist)

# create null networks
set.seed(1)
net_layout<-ggnetwork(net,layout=with_fr())
set.seed(3)
net_rand<-sample_fitness(ecount(net),degree(net))
vertex_attr(net_rand,"name")<-V(net)$name
edge_attr(net_rand)<-apply(get.edgelist(net_rand),1,function(x) {
  ifelse(x[1]>x[2],paste(x[2],x[1],sep="-"),paste(x[1],x[2],sep="-"))}) %>%
  data.frame(pair=.,stringsAsFactors=F) %>%
  left_join(pair_dist) %>%
  rename(pairwise=dist) %>%
  left_join(phylo_dist) %>%
  rename(phylo=dist) %>%
  as.list
net_rand_layout<-data.frame(get.edge.attribute(net_rand)) %>%
  separate(pair,c("from","to"),sep="-") %>%
  full_join(unique(select(net_layout,x,y,name)),by=c("to"="name")) %>%
  rename(xend=x,yend=y) %>%
  full_join(unique(select(net_layout,x,y,name)),by=c("from"="name")) %>%
  mutate(x=ifelse(is.na(x),xend,x),y=ifelse(is.na(y),yend,y),
         xend=ifelse(is.na(xend),x,xend),yend=ifelse(is.na(yend),y,yend),
         from=ifelse(is.na(from),to,from),to=ifelse(is.na(to),from,to)) %>%
  unique()

set.seed(1)
net_full_layout<-rbind(rbind(mutate(net_layout,method="observed"),
                       mutate(net_layout,method="shuffled tree tips",
                              pairwise=ifelse(is.na(pairwise),NA,sample(pairwise[!is.na(pairwise)])))) %>%
                         select(-cor),
                       mutate(net_rand_layout,method="random network") %>%
                         select(-to) %>% rename(name=from)) %>%
  mutate(method=factor(method,levels=unique(method)))

plot_net<-ggplot(net_full_layout,aes(x=x,y=y,xend=xend,yend=yend)) +
  geom_edges(aes(color=pairwise)) +
  geom_nodes(size=2,color="grey50") +
  scale_color_gradientn(colors = hue_pal()(5),values=rescale(quantile(seq(0,50))/100)) +
  facet_wrap(~method) +
  # expand_limits(y=-0.05) +
  theme_blank(strip.text=element_text(face="bold"),
              strip.background=element_rect(color=NA),
              legend.direction="horizontal",
              legend.position=c(0.67,0.065),
              legend.title=element_text(size=9),
              legend.text=element_text(size=8),
              plot.margin=margin(3,3,3,3,"pt"),
              legend.margin=margin(0,0,0,0,"pt"),
              legend.key.height=unit(0.7,"line"),
              legend.key.width=unit(1.5,"line")) +
  labs(color="sequence dissimilarity ") +
  guides(color=guide_colorbar(title.position="top",title.hjust=0.5))
ggsave("Figure_1.pdf",plot_net,width=10,height=3.5)

##########################
### network statistics ###
##########################
# nb samples, threshold, nb candidate OTUs, nb otu (%), nb candidate edges, nb edges (%), average degree, average path length
net_stat<-ddply(files,.(project,type),function(i){
  tmp_net<-get(paste0(i$file,"_net"))
  tmp_cand<-get(paste0(as.character(i$project),"_mat"))
  tmp_mat<-tmp_cand[,V(tmp_net)$name]
  data.frame(samples=nrow(tmp_cand),
             cand_otu=ncol(tmp_cand),
             cand_reads=sum(tmp_cand),
             threshold=scan(file.path(paste0("NetworkNullHPC.",i$nnhpc),ifelse(i$type=="cooc","threshold","ex_threshold")),quiet=T),
             net_otu=ncol(tmp_mat),
             net_reads=sum(tmp_mat),
             cand_edges=ncol(tmp_cand)*(ncol(tmp_cand)-1)/2,
             net_edges=ecount(tmp_net),
             average_degree=mean(degree(tmp_net)),
             average_path_length=average.path.length(tmp_net))
}) %>%
  mutate(project=mapvalues(project,levels(project),c("Neotropical soil","marine DCM","marine surface")),
         project=factor(project,rev(levels(project))),
         type=mapvalues(type,levels(type),c("co-exclusion","co-occurrence")),
         type=factor(type,rev(levels(type))),
         otu_perc=net_otu/cand_otu*100,edge_perc=net_edges/cand_edges*100) %>%
  arrange(type,project)
write.table(net_stat,"Table_1.tsv",row.names=F,quote=F,sep="\t")


#########################
## Occurrence profiles ##
#########################
oc<-foreach(i=isplit(files,files$project),.combine=rbind) %do% {
  tmp_mat<-get(paste0(i$key[[1]],"_mat"))
  tmp_env<-get(paste0(i$key[[1]],"_env"))
  rbind(data.frame(tmp_mat) %>%
          rownames_to_column("sample") %>%
          gather("OTU","reads",-sample) %>%
          filter(reads>0) %>%
          left_join(tmp_env) %>%
          group_by(OTU,unit) %>%
          summarize(oc=n()) %>%
          group_by(OTU) %>%
          summarize(units=n(),occurrence=sum(oc)) %>%
          mutate(network="cand"),
        foreach(j=isplit(i$value,i$value$type),.combine=rbind) %do% {
          tmp_net<-get(paste(i$key[[1]],j$key[[1]],"net",sep="_"))
          tmp_net_mat<-tmp_mat[,V(tmp_net)$name]
          data.frame(tmp_net_mat) %>%
            rownames_to_column("sample") %>%
            gather("OTU","reads",-sample) %>%
            filter(reads>0) %>%
            left_join(tmp_env) %>%
            group_by(OTU,unit) %>%
            summarize(oc=n()) %>%
            group_by(OTU) %>%
            summarize(units=n(),occurrence=sum(oc)) %>%
            mutate(network=j$key[[1]])
        }) %>%
    mutate(project=i$key[[1]])
} %>%
  mutate(network=as.factor(network),
         network=fct_relevel(network,"coex",after=2),
         network=mapvalues(network,levels(network),c("candidate OTUs","co-occurring OTUs","co-excluding OTUs")),
         project=fct_relevel(as.factor(project),rev),
         project=mapvalues(project,levels(project),c("marine surface","marine DCM","Neotropical soil"))) %>%
  group_by(project,network,occurrence,units) %>%
  summarize(count=n()) %>%
  ungroup() %>%
  mutate(units=factor(units))

plot_oc<-dlply(oc,.(project),function(x) {
  ggplot(x,aes(occurrence,count,fill=units)) +
    geom_area(position="stack") +
    scale_fill_discrete(h=c(0,300),drop=F) +
    facet_wrap(~network,scales="free_y") +
    labs(fill="geographical units: ") +
    theme(legend.position="bottom",
          legend.margin=margin(0,1,0,1,"line"),
          legend.key.width=unit(1.2,"line"),
          legend.background=element_blank()) +
    guides(fill=guide_legend(nrow=1))
})
grid_oc<-ggarrange(plot_oc[[1]] + theme(axis.title.x=element_blank()) + guides(fill=F),
                   plot_oc[[2]] + theme(axis.title.x=element_blank()) + guides(fill=F),
                   plot_oc[[3]],
                   labels=c("A","B","C"),heights=c(0.8,0.8,1),ncol=1)
ggsave("Figure_S2.pdf",grid_oc,width=8,height=8)


##############
## taxonomy ##
##############
taxo<-foreach(i=isplit(files,files$project),.combine=rbind) %do% {
  tmp_taxo<-get(paste0(i$key[[1]],"_taxo")) %>%
    mutate(taxo=ifelse(Phylum=="Stramenopiles_X",as.character(Class),as.character(Phylum)),
           total=total/sum(total),
           oc=1/n()) %>%
    select(OTU,taxo,total,oc)
  foreach(j=isplit(i$value,i$value$type),.combine=rbind) %do% {
    tmp_net<-get(paste(i$key[[1]],j$key[[1]],"net",sep="_"))
    filter(tmp_taxo,OTU %in% V(tmp_net)$name) %>%
      mutate(type=j$key[[1]])
  } %>%
    rbind(mutate(tmp_taxo,type="cand"),.) %>%
    group_by(type,taxo) %>%
    summarize(reads=sum(total),OTUs=sum(oc)) %>%
    mutate(project=i$key[[1]])
} %>% ungroup() %>%
  mutate(taxo=ifelse(is.na(taxo),"unidentified",taxo),
         taxo=as.factor(taxo),
         type=as.factor(type),
         type=fct_relevel(type,"coex",after=2),
         type=mapvalues(type,levels(type),c("candidate OTUs","co-occurring OTUs","co-excluding OTUs")),
         project=fct_relevel(as.factor(project),rev),
         project=mapvalues(project,levels(project),c("marine surface","marine DCM","Neotropical soil")))

# select the most abundant and OTU rich taxa: average relative abundance and number of OTU of at least 1 % 
taxa<-filter(taxo,type=="candidate OTUs",reads>0.01,OTUs>0.01) %>%
  group_by(taxo) %>%
  summarize(reads=mean(reads)) %>%
  arrange(reads)
taxo_df<-mutate(taxo,taxo=ifelse(taxo %in% taxa$taxo,as.character(taxo),"other protists"),
                taxo=factor(taxo,levels=c("other protists",as.character(taxa$taxo))),
                taxo=mapvalues(taxo,levels(taxo),sub("-","-\n",levels(taxo)))) %>%
  group_by(project,type,taxo) %>%
  summarize(reads=sum(reads),OTUs=sum(OTUs)) %>%
  gather("method","value",reads,OTUs)

interleave <- function(v1,v2,rev=F) {
  if(missing(v2)) {
    ord1 <- seq(1,length(v1),2)
    ord2 <- seq(2,length(v1),2)
    v1[order(c(ord1,ord2))]
  } else {
    ord1 <- 2*(1:length(v1))-1
    ord2 <- 2*(1:length(v2))
    c(v1,v2)[order(c(ord1,ord2))]
  }  
}
taxo_colors<-rev(interleave(diag(as.matrix(ldply(interleave(seq(50,80,length.out=length(levels(taxo_df$taxo)))),
                                                 function(x) hue_pal(h=c(10,300),l=x)(length(levels(taxo_df$taxo))))))))

plot_taxo<-ggplot(taxo_df,aes(type,value,fill=taxo)) +
  geom_bar(stat="identity",position="stack") +
  facet_grid(method~project) +
  scale_y_continuous(labels=scales::percent) +
  scale_fill_manual(values=taxo_colors) +
  labs(y="percents",fill="taxonomy") +
  theme(axis.ticks.x = element_blank(),
        axis.text.x=element_text(angle=60,hjust=1.1,vjust=1.1),
        axis.title.x = element_blank(),
        legend.background=element_blank(),
        legend.title=element_text(face="bold")) +
  guides(fill=guide_legend(ncol=1))
ggsave("Figure_S3.pdf",plot_taxo,width=6.5,height=8)

#########
## SES ##
#########
# read all ses results to a single data-frame
ses<-ddply(files,.(project,type),function(x) {
  readRDS(paste0(x$file,"_ses_df"))
}) %>%
  mutate(project=mapvalues(project,levels(project),c("Neotropical soil","marine DCM","marine surface")),
         project=fct_relevel(project,rev),
         type=mapvalues(type,levels(type),c("co-exclusion","co-occurrence")),
         type=fct_relevel(type,rev),
         signif=factor(signif,levels=c("TRUE","FALSE")),
         method=mapvalues(method,levels(method),c("Sequence based","Phylogenetic based")))
ses2<-mutate(ses,significance=ifelse(signif==F,"n.s.",ifelse(pval<=0.001,"***",ifelse(pval<=0.01,"**","*"))),
             significance=factor(significance,levels=c("***","**","*","n.s.")))

# color continuous distance area of simultaneous co-occurrence and co-exclusion SES directions
area_color_groups<-mutate(ses2,effect=ifelse(signif==T,ifelse(ses>0,"positive","negative"),"null")) %>%
  select(project,type,method,dist,model,effect) %>%
  spread(type,effect) %>%
  rename(direction=`co-occurrence`) %>%
  mutate(direction=ifelse(direction==`co-exclusion`,direction,"null")) %>%
  group_by(model,project,method,direction) %>%
  arrange(model,project,method,direction,dist) %>%
  mutate(diff=as.numeric(as.character(mapvalues(method,levels(method),c(0.015,0.15)))),
         subgroup = cumsum((dist - lag(dist)) > diff & !is.na(lag(dist)))) %>%
  group_by(model,project,method,direction,subgroup) %>%
  summarize(xmin=min(dist),xmax=max(dist)) %>%
  mutate(xmin=ifelse(method=="Phylogenetic based",xmin-0.05,xmin-0.005),
         xmax=ifelse(method=="Phylogenetic based",xmax+0.05,xmax+0.005)) %>%
  filter(direction!="null") %>%
  droplevels()

plot_ses_tree2<-ggplot(filter(ses2,model=="shuffled tree tips"),aes(dist,ses,color=type,fill=type)) +
  geom_rect(data=filter(area_color_groups,model=="shuffled tree tips",direction=="positive"),
            aes(xmin=xmin,xmax=xmax),ymin=-Inf,ymax=Inf,
            alpha=0.4,inherit.aes=F,fill="#0072B2") +
  geom_rect(data=filter(area_color_groups,model=="shuffled tree tips",direction=="negative"),
            aes(xmin=xmin,xmax=xmax),ymin=-Inf,ymax=Inf,
            alpha=0.2,inherit.aes=F,fill="#CC79A7") +
  geom_smooth(method="loess",se=F,span=0.5,size=0.7) +
  geom_point(aes(shape=significance,size=significance),alpha=0.3) +
  scale_shape_manual(values=c(21,24,23,4)) +
  scale_size_manual(values=c(rep(1.8,3),1.2)) +
  # scale_alpha_manual(values=c(rep(0.7,3),0.8)) +
  scale_color_manual(values=rev(colorblind_pal()(4)[c(2,4)])) +
  scale_fill_manual(values=rev(colorblind_pal()(4)[c(2,4)])) +
  facet_grid(project~method,scales="free") +
  labs(x="distance",y="Standardize effect size",color="network",fill="network") +
  theme(legend.position=c(0.87,0.14))
gt_ses_tree2<-ggplot_gtable(ggplot_build(plot_ses_tree2))
gt_ses_tree2$widths[7]<-1.5*gt_ses_tree2$widths[7]
gt_ses_tree2$heights[12]<-0.8*gt_ses_tree2$heights[12]
ggsave("Figure_2.pdf",gt_ses_tree2,width=7,height=9)

plot_ses_net2<-ggplot(filter(ses2,model=="random network"),aes(dist,ses,color=type,fill=type)) +
  geom_rect(data=filter(area_color_groups,model=="random network",direction=="positive"),
            aes(xmin=xmin,xmax=xmax),ymin=-Inf,ymax=Inf,
            alpha=0.4,inherit.aes=F,fill="#0072B2") +
  geom_rect(data=filter(area_color_groups,model=="random network",direction=="negative"),
            aes(xmin=xmin,xmax=xmax),ymin=-Inf,ymax=Inf,
            alpha=0.2,inherit.aes=F,fill="#CC79A7") +
  geom_smooth(method="loess",se=F,span=0.5,size=0.7) +
  geom_point(aes(shape=significance,size=significance),alpha=0.3) +
  scale_shape_manual(values=c(21,24,23,4)) +
  scale_size_manual(values=c(rep(1.8,3),1.2)) +
  # scale_alpha_manual(values=c(rep(0.7,3),0.8)) +
  scale_color_manual(values=rev(colorblind_pal()(4)[c(2,4)])) +
  scale_fill_manual(values=rev(colorblind_pal()(4)[c(2,4)])) +
  facet_grid(project~method,scales="free") +
  labs(x="distance",y="Standardize effect size",color="network",fill="network") +
  theme(legend.position=c(0.87,0.14))
gt_ses_net2<-ggplot_gtable(ggplot_build(plot_ses_net2))
gt_ses_net2$widths[7]<-1.5*gt_ses_net2$widths[7]
gt_ses_net2$heights[12]<-0.8*gt_ses_net2$heights[12]
ggsave("Figure_S4.pdf",gt_ses_net2,width=7,height=9)

######################
## network coverage ##
######################
coverage<-ddply(files,.(project,type),function(x) {
  readRDS(paste0(x$file,"_network_coverage"))
}) %>% mutate(project=mapvalues(project,levels(project),c("Neotropical soil","marine DCM","marine surface")),
              project=factor(project,rev(levels(project))),
              type=mapvalues(type,levels(type),c("co-exclusion","co-occurrence")),
              type=factor(type,rev(levels(type))),
              method=mapvalues(method,levels(method),c("Sequence based","Phylogenetic based")),
              dist=as.numeric(as.character(dist)))

plot_cov<-ggplot(coverage,aes(dist,percent)) +
  geom_line(aes(color=project,linetype=type)) +
  facet_wrap(~method,scales="free") +
  scale_color_manual(values=colorblind_pal()(7)[c(4,7,3)]) +
  scale_y_log10() +
  labs(x="distance",color="Dataset:",linetype="Network:") +
  theme(legend.position="bottom",
        legend.background=element_blank(),
        legend.spacing.x=unit(0.3,"line"),
        legend.margin=margin(0,1,0,1,"line"),
        legend.key.width=unit(1.2,"line"),
        plot.margin=margin(5,5,2,5,"pt")) +
  guides(linetype=guide_legend(override.aes=list(size=1)),
         color=guide_legend(override.aes=list(size=1)))
ggsave("Figure_S5.pdf",plot_cov,width=8,height=3.3)


##############################
## shared taxonomy of edges ##
##############################
# reduce taxonomy to the correctly classified levels
for(i in levels(files$project)) {
  tmp_agg<-get(paste0(i,"_taxo")) %>%
    select(matches("^[A-Z][A-Za-z]*$",ignore.case=F)) %>%
    unite("taxo",-OTU,sep="|") %>%
    mutate(taxo=sub("(unclassified\\|)\\1*","",sub("\\|([^\\|]*)\\|.*\\1[_X]*\\+sp\\.\\|$","\\|\\1\\|",sub("$","\\|",gsub("\\|NA","",taxo)))))
  assign(paste0(i,"_taxo_agg"),tmp_agg)
  write.table(tmp_agg,paste0(i,"_taxo_agg.tsv"),col.names=F,row.names=F,sep="\t",quote=F)
}

# shared taxonomy of observed edges (avoid _X levels: go to the deepest correctly classified shared taxa)
pair_taxo<-foreach(i=isplit(files,files$project),.combine=rbind) %do% {
  tmp_taxo<-get(paste0(i$key[[1]],"_taxo_agg"))
  foreach(j=isplit(i$value,i$value$type),.combine=rbind) %do% {
    get.edgelist(get(paste(i$key[[1]],j$key[[1]],"net",sep="_"))) %>%
      data.frame(stringsAsFactors=F)  %>%
      setNames(c("from","to")) %>%
      left_join(tmp_taxo,by=c("from"="OTU")) %>%
      rename(taxo1=taxo) %>%
      left_join(tmp_taxo,by=c("to"="OTU")) %>%
      unite("both",taxo1,taxo,sep="#",remove=T) %>%
      mutate(common=sub("\\|([^\\|]*)\\|.*\\1[_X]*$","\\|\\1",sub("^(.*)\\|.*#\\1\\|.*","\\1",both))) %>%
      group_by(common) %>%
      summarize(n=n()) %>%
      mutate(percent=n/sum(n)*100) %>%
      arrange(desc(n)) %>%
      mutate(project=i$key[[1]],type=j$key[[1]]) 
  }
}
pair_level<-adply(pair_taxo,1,function(x) {
  data.frame(x,level=length(unlist(strsplit(x$common,"\\|"))))}) %>%
  group_by(project,type,level) %>%
  summarize(obs=sum(n),obs_percent=sum(percent)) %>%
  arrange(project,type,level)

# compare to all candidate edges
all_pair_taxo<-foreach(i=isplit(files,files$project),.combine=rbind) %do% {
  tmp_taxo<-get(paste0(i$key[[1]],"_taxo_agg"))
  foreach(j=isplit(i$value,i$value$type),.combine=rbind) %do% {
    tmp_net<-get(paste(i$key[[1]],j$key[[1]],"net",sep="_"))
    setNames(data.frame(t(combn(V(tmp_net)$name,2))),c("from","to")) %>%
      left_join(tmp_taxo,by=c("from"="OTU")) %>%
      rename(taxo1=taxo) %>%
      left_join(tmp_taxo,by=c("to"="OTU")) %>%
      unite("both",taxo1,taxo,sep="#",remove=T) %>%
      mutate(common=sub("\\|([^\\|]*)\\|.*\\1[_X]*$","\\|\\1",sub("^(.*)\\|.*#\\1\\|.*","\\1",both))) %>%
      group_by(common) %>%
      summarize(n=n()) %>%
      mutate(percent=n/sum(n)*100) %>%
      arrange(desc(n)) %>%
      mutate(project=i$key[[1]],type=j$key[[1]])
  }
}

all_pair_level<-adply(all_pair_taxo,1,function(x) {data.frame(x,level=length(unlist(strsplit(x$common,"\\|"))))}) %>%
  group_by(project,type,level) %>%
  summarize(total=sum(n),total_percent=sum(percent)) %>%
  ungroup() %>%
  arrange(project,type,level) %>%
  left_join(pair_level) %>%
  replace_na(list(obs=0,obs_percent=0)) %>%
  mutate(percent=obs/total*100,
         level=factor(as.character(level),as.character(1:8)),
         level=mapvalues(level,levels(level),grep("^[A-Z][a-z]*$",colnames(tara_SUR_taxo),value="T")),
         type=as.factor(type),
         type=fct_relevel(type,"coex",after=1),
         type=mapvalues(type,levels(type),c("co-occurrence","co-exclusion")),
         project=as.factor(project),
         project=fct_relevel(project,rev),
         project=mapvalues(project,levels(project),c("marine surface","marine DCM","Neotropical soil"))) %>%
  complete(crossing(project,type,level),fill=list(percent=0))

# plot
plot_edge_taxo<-ggplot(all_pair_level,aes(level,percent,fill=type)) +
  geom_bar(stat="identity",position="dodge") +
  facet_wrap(~project,scales="free_y",ncol=1) +
  scale_fill_manual(values=colorblind_pal()(4)[c(4,2)]) +
  # scale_y_sqrt() +
  labs(x="shared taxonomic levels", fill="network",y="percent of candidate edges") +
  theme(legend.position=c(0.2,0.57))
ggsave("Figure_S6.pdf",plot_edge_taxo,width=4.5,height=8)


####################################
## shared taxonomy along distance ##
####################################
# observed networks
edge_taxo_dist<-foreach(i=isplit(files,files$project),.combine=rbind) %do% {
  tmp_taxo<-get(paste0(i$key[[1]],"_taxo_agg"))
  foreach(j=isplit(i$value,i$value$type),.combine=rbind) %do% {
    get.edgelist(get(paste(i$key[[1]],j$key[[1]],"net",sep="_"))) %>%
      data.frame(stringsAsFactors=F)  %>%
      setNames(c("from","to")) %>%
      cbind.data.frame(get.edge.attribute(get(paste(i$key[[1]],j$key[[1]],"net",sep="_")))) %>%
      left_join(tmp_taxo,by=c("from"="OTU")) %>%
      rename(taxo1=taxo) %>%
      left_join(tmp_taxo,by=c("to"="OTU"))  %>%
      mutate(project=i$key[[1]],type=j$key[[1]]) %>%
      unite("both",taxo1,taxo,sep="#",remove=T) %>%
      mutate(common=sub("\\|([^\\|]*)\\|.*\\1[_X]*$","\\|\\1",sub("^(.*)\\|.*#\\1\\|.*","\\1",both))) %>%
      rowwise() %>%
      mutate(level=length(unlist(strsplit(common,"\\|"))))
  }
}
edge_taxo_dist_stat<-gather(edge_taxo_dist,"method","dist",pairwise,phylo) %>%
  group_by(method) %>%
  mutate(digits=ifelse(method=="phylo",1,2),
         max=ifelse(method=="phylo",9.9,0.49),
         dist=ifelse(dist>max,max,floor(dist*10^digits)*10^-digits)) %>%
  group_by(project,type,method,dist,level) %>%
  summarize(n=n()) %>%
  mutate(percent=n/sum(n)*100) %>%
  ungroup() %>%
  mutate(dist=ifelse(method=="phylo",dist+0.05,dist+0.005),
         level=factor(as.character(level),as.character(1:8)),
         level=mapvalues(level,levels(level),grep("^[A-Z][a-z]*$",colnames(tara_SUR_taxo),value="T")),
         type=as.factor(type),
         type=fct_relevel(type,"coex",after=1),
         type=mapvalues(type,levels(type),c("co-occurrence","co-exclusion")),
         project=as.factor(project),
         project=fct_relevel(project,rev),
         project=mapvalues(project,levels(project),c("marine surface","marine DCM","Neotropical soil"))) %>%
  arrange(method,project,type,dist,level)

# test if more taxonomic depth is covered in co-occurrence compared to co-exclusion networks for the same distance class
edge_taxo_dist_test<-mutate(edge_taxo_dist_stat,level=as.numeric(as.character(mapvalues(level,levels(level),1:8)))) %>%
  select(-percent) %>%
  ddply(.(project,method,dist),function(x) {
    if(length(unique(x$type))==2) {
      ldply(c("greater","less"),function(y) {
        c(dir=y,pval=wilcox.test(level~type,data=uncount(x,n),alternative=y)$p.value)
      })
    } else {
      return(data.frame(dir=NA,pval=NA))
    }
  }) %>%
  filter(!is.na(pval)) %>%
  mutate(signif=pval<=0.05)

## shared taxonomic levels of all candidate edges for each distance class ##
all_edges_taxo<-foreach(i=levels(files$project),.combine=rbind) %do% {
  tmp_taxo<-get(paste0(i,"_taxo_agg"))
  tmp_tree<-read.tree(file.path("phylogeny",i,paste0("RAxML_bestTree.MultipleOriginal_",i)))
  tmp_dist<-read.table(paste0(i,".suma.pairwise"),col.names=c("from","to","pairwise"),stringsAsFactors=F) %>%
    mutate(pair=paste(from,to,sep="-")) %>%
    select(pair,pairwise) %>%
    left_join(setNames(data.frame(t(combn(tmp_tree$tip.label,2)),c(as.dist(cophenetic(tmp_tree))),
                                  stringsAsFactors=F),
                       c("from","to","phylo")) %>%
                mutate(pair=ifelse(from<to,paste(from,to,sep="-"),paste(to,from,sep="-"))) %>%
                select(pair,phylo)) %>%
    separate(pair,c("from","to"),sep="-") %>%
    left_join(tmp_taxo,by=c("from"="OTU")) %>%
    rename(taxo1=taxo) %>%
    left_join(tmp_taxo,by=c("to"="OTU")) %>%
    unite("both",taxo1,taxo,sep="#",remove=T) %>%
    mutate(common=sub("\\|([^\\|]*)\\|.*\\1[_X]*$","\\|\\1",sub("^(.*)\\|.*#\\1\\|.*","\\1",both)),
           pairwise=ifelse(pairwise>0.5,0.5,floor(pairwise*100)/100),
           phylo=ifelse(phylo>10,10,floor(phylo*10)/10)) %>%
    gather("method","dist",pairwise,phylo) %>%
    select(-both)
  return(foreach(j=levels(files$type),.combine=rbind) %do% {
    tmp_nodes<-V(get(paste(i,j,"net",sep="_")))$name
    filter(tmp_dist,from %in% tmp_nodes,to %in% tmp_nodes) %>%
      mutate(type=j)
  } %>%
    group_by(type,method,dist,common) %>%
    summarize(n=n()) %>%
    mutate(percent=n/sum(n)*100) %>%
    arrange(desc(n)) %>%
    mutate(project=i) %>%
    data.frame())
  rm(list=ls(pattern="^tmp"))
}

all_edges_level<-adply(all_edges_taxo,1,function(x) {data.frame(x,level=length(unlist(strsplit(x$common,"\\|"))))}) %>%
  group_by(project,method,type,dist,level) %>%
  summarize(total=sum(n),total_percent=sum(percent)) %>%
  ungroup() %>%
  mutate(level=factor(as.character(level),as.character(1:8)),
         level=mapvalues(level,levels(level),grep("^[A-Z][a-z]*$",colnames(tara_SUR_taxo),value="T")),
         type=as.factor(type),
         type=fct_relevel(type,"coex",after=1),
         type=mapvalues(type,levels(type),c("co-occurrence","co-exclusion")),
         project=as.factor(project),
         project=fct_relevel(project,rev),
         project=mapvalues(project,levels(project),c("marine surface","marine DCM","Neotropical soil")),
         dist=ifelse(method=="phylo",dist+0.05,dist+0.005))%>%
  arrange(project,method,type,dist,level)

# test if more taxonomic depth is covered in all candidate edges of the co-occurrence compared to the co-exclusion networks for the same distance class
all_edges_test<-mutate(all_edges_level,level=as.numeric(as.character(mapvalues(level,levels(level),1:8)))) %>%
  select(-total_percent) %>%
  uncount(total) %>%
  ddply(.(project,method,dist),function(x) {
    if(length(unique(x$type))==2) {
      ldply(c("greater","less"),function(y) {
        c(dir=y,pval=wilcox.test(level~type,data=x,alternative=y)$p.value)
      })
    } else {
      return(data.frame(dir=NA,pval=NA))
    }
  }) %>%
  filter(!is.na(pval)) %>%
  mutate(signif=pval<=0.05,
         y=mapvalues(dir,c("greater","less"),c(-1,101)))

# test if more/less taxonomic depth is covered in observed networks compared to candidate edges for the same distance class
obs_vs_all_edges_test<-left_join(all_edges_level,edge_taxo_dist_stat) %>%
  select(-percent,-total_percent) %>%
  drop_na(n) %>%
  gather("measure","count",n,total,factor_key=T) %>%
  mutate(level=as.numeric(as.character(mapvalues(level,levels(level),1:8)))) %>%
  ddply(.(project,type,method,dist),function(x) {
    if(nrow(filter(x,measure=="n"))>0) {
      ldply(c("greater","less"),function(y) {
        c(dir=y,pval=wilcox.test(level~measure,data=uncount(x,count),alternative=y)$p.value)
      })
    } else {
      return(data.frame(dir=NA,pval=NA))
    }
  }) %>%
  filter(!is.na(pval)) %>%
  mutate(signif=pval<=0.05,
         y=ifelse(dir=="less",102,-2))

# figures
plot_edge_taxo_pairwise<-ggplot(filter(edge_taxo_dist_stat,method=="pairwise"),aes(dist,percent)) +
  geom_rect(data=filter(area_color_groups,model=="shuffled tree tips",direction=="positive",method=="Sequence based"),
            aes(xmin=xmin,xmax=xmax),ymin=-Inf,ymax=Inf,
            alpha=0.4,inherit.aes=F,fill="#0072B2") +
  geom_rect(data=filter(area_color_groups,model=="shuffled tree tips",direction=="negative",method=="Sequence based"),
            aes(xmin=xmin,xmax=xmax),ymin=-Inf,ymax=Inf,
            alpha=0.2,inherit.aes=F,fill="#CC79A7") +
  geom_point(data=filter(obs_vs_all_edges_test,method=="pairwise",signif==T),
             aes(x=dist,y=y),shape=8,size=0.5,inherit.aes=F) +
  geom_bar(aes(fill=level),position="stack",stat="identity",show.legend=F) +
  facet_grid(project~type) +
  scale_fill_brewer(palette="Accent") +
  labs(x="pairwise dissimilarity")
gt_edge_taxo_pairwise<-ggplotGrob(plot_edge_taxo_pairwise)
plot_edge_taxo_phylo<-ggplot(filter(edge_taxo_dist_stat,method=="phylo"),aes(dist,percent,fill=level)) +
  geom_rect(data=filter(area_color_groups,model=="shuffled tree tips",direction=="positive",method=="Phylogenetic based"),
            aes(xmin=xmin,xmax=xmax),ymin=-Inf,ymax=Inf,
            alpha=0.4,inherit.aes=F,fill="#0072B2") +
  geom_rect(data=filter(area_color_groups,model=="shuffled tree tips",direction=="negative",method=="Phylogenetic based"),
            aes(xmin=xmin,xmax=xmax),ymin=-Inf,ymax=Inf,
            alpha=0.2,inherit.aes=F,fill="#CC79A7") +
  geom_point(data=filter(obs_vs_all_edges_test,method=="phylo",signif==T),
             aes(x=dist,y=y),shape=8,size=0.5,inherit.aes=F) +
  geom_bar(position="stack",stat="identity") +
  facet_grid(project~type) +
  scale_fill_brewer(palette="Accent") +
  labs(x="phylogenetic distance",fill="PR² ranks") +
  theme(legend.title=element_text(hjust=0.5),
        legend.background=element_blank(),
        legend.position=c(0.85,0.16),
        plot.margin=margin(5.5,5.5,5.5,20,"pt"))
gt_edge_taxo_phylo<-ggplotGrob(plot_edge_taxo_phylo)
plot_edge_taxo_dist<-ggarrange(gt_edge_taxo_pairwise[,-8],gt_edge_taxo_phylo[,-c(3,4)],labels=c("A","B"),ncol=2,widths=c(0.8,1))
ggsave("Figure_3.pdf",plot_edge_taxo_dist,width=9,height=7)

plot_all_edges_level_pairwise<-ggplot(filter(all_edges_level,method=="pairwise"),aes(dist,total_percent)) +
  geom_rect(data=filter(area_color_groups,model=="shuffled tree tips",direction=="positive",method=="Sequence based"),
            aes(xmin=xmin,xmax=xmax),ymin=-Inf,ymax=Inf,
            alpha=0.4,inherit.aes=F,fill="#0072B2") +
  geom_rect(data=filter(area_color_groups,model=="shuffled tree tips",direction=="negative",method=="Sequence based"),
            aes(xmin=xmin,xmax=xmax),ymin=-Inf,ymax=Inf,
            alpha=0.2,inherit.aes=F,fill="#CC79A7") +
  geom_bar(aes(fill=level),position="stack",stat="identity",show.legend=F) +
  facet_grid(project~type) +
  scale_fill_brewer(palette="Accent") +
  labs(x="pairwise dissimilarity",y="percent")
gt_all_edges_level_pairwise<-ggplotGrob(plot_all_edges_level_pairwise)
plot_all_edges_level_phylo<-ggplot(filter(all_edges_level,method=="phylo"),aes(dist,total_percent,fill=level)) +
  geom_rect(data=filter(area_color_groups,model=="shuffled tree tips",direction=="positive",method=="Phylogenetic based"),
            aes(xmin=xmin,xmax=xmax),ymin=-Inf,ymax=Inf,
            alpha=0.4,inherit.aes=F,fill="#0072B2") +
  geom_rect(data=filter(area_color_groups,model=="shuffled tree tips",direction=="negative",method=="Phylogenetic based"),
            aes(xmin=xmin,xmax=xmax),ymin=-Inf,ymax=Inf,
            alpha=0.2,inherit.aes=F,fill="#CC79A7") +
  geom_bar(position="stack",stat="identity") +
  facet_grid(project~type) +
  scale_fill_brewer(palette="Accent") +
  labs(x="phylogenetic distance",fill="PR² ranks") +
  theme(legend.title=element_text(hjust=0.5),
        legend.background=element_blank(),
        legend.position=c(0.85,0.16),
        plot.margin=margin(5.5,5.5,5.5,20,"pt"))
gt_all_edges_level_phylo<-ggplotGrob(plot_all_edges_level_phylo)
plot_all_edges_level<-ggarrange(gt_all_edges_level_pairwise[,-8],gt_all_edges_level_phylo[,-c(3,4)],labels=c("A","B"),ncol=2,widths=c(0.8,1))
ggsave("Figure_S7.pdf",plot_all_edges_level,width=9,height=7)


####################################
## taxonomy of the inversion zone ##
####################################
ocean<-filter(files,grepl("tara",project)) %>%
  droplevels()
invz<-filter(area_color_groups,direction=="positive",method=="Sequence based")
system(paste0("xmin=",invz$xmin,"; xmax=",invz$xmax,
              "; awk -v x=$xmin -v X=$xmax '$3>=x && $3<X{print}' tara_SUR.suma.pairwise > tara_SUR.suma.pairwise.invz"))
system(paste0("xmin=",invz$xmin,"; xmax=",invz$xmax,
              "; awk -v x=$xmin -v X=$xmax '$3>=x && $3<X{print}' tara_DCM.suma.pairwise > tara_DCM.suma.pairwise.invz"))

invz_taxo<-foreach(i=isplit(ocean,ocean$project),.combine=rbind) %do% {
  tmp_agg<-get(paste0(i$key[[1]],"_taxo_agg"))
  foreach(j=isplit(i$value,i$value$type),.combine=rbind) %do% {
    data.frame(i$key[[1]],j$key[[1]],get.edgelist(get(paste(i$key[[1]],j$key[[1]],"net",sep="_"))),stringsAsFactors=F) %>%
      setNames(c("project","type","from","to"))
  } %>%
    inner_join(read.table(paste0(i$key[[1]],".suma.pairwise.invz"),col.names=c("from","to","dist"),stringsAsFactors=F)) %>%
    left_join(get(paste0(i$key[[1]],"_taxo_agg")),by=c("from"="OTU")) %>%
    rename(taxo1=taxo) %>%
    left_join(get(paste0(i$key[[1]],"_taxo_agg")),by=c("to"="OTU"))
}
invz_pair_taxo<-unite(invz_taxo,"both",taxo1,taxo,sep="#",remove=T) %>%
  mutate(common=sub("^(.*)\\|.*#\\1\\|.*","\\1",both)) %>%
  group_by(project,type,common) %>%
  summarize(n=n()) %>%
  mutate(percent=n/sum(n)*100)
  
invz_level<-adply(invz_pair_taxo,1,function(x) {data.frame(x,level=length(unlist(strsplit(x$common,"\\|"))))}) %>%
  group_by(project,type,level) %>%
  summarize(n=sum(n),percent=sum(percent)) %>%
  arrange(project,type,level) %>%
  left_join(pair_level) %>%
  mutate(ratio=n/obs*100)

# only for Surface
invz_taxo_class<-filter(invz_taxo,project=="tara_SUR") %>%
  select(-project,-from,-to) %>%
  mutate(taxo1=ifelse(grepl("Chrysophyceae-Synurophyceae|MAST|MOCH|Bacillariophyta",taxo1),
                      sub("^(([^\\|]*\\|){4}).*$","\\1",taxo1,perl=T),
                      sub("^(([^\\|]*\\|){3}).*$","\\1",taxo1,perl=T)),
         taxo=ifelse(grepl("Chrysophyceae-Synurophyceae|MAST|MOCH|Bacillariophyta",taxo),
                     sub("^(([^\\|]*\\|){4}).*$","\\1",taxo,perl=T),
                     sub("^(([^\\|]*\\|){3}).*$","\\1",taxo,perl=T)),
         taxo1=sub("Alveolata","other_Alveolata",sub("Stramenopiles","other_Stramenopiles",sub("_X$","",sub("^.*\\|([^\\|]*)\\|$","\\1",taxo1,perl=T)))),
         taxo=sub("Alveolata","other_Alveolata",sub("Stramenopiles","other_Stramenopiles",sub("_X$","",sub("^.*\\|([^\\|]*)\\|$","\\1",taxo,perl=T))))) %>%
  rowwise() %>%
  mutate(from=sort(c(taxo1,taxo))[1],
         to=sort(c(taxo1,taxo))[2]) %>%
  group_by(type,from,to) %>%
  summarize(n=n()) %>%
  ungroup() %>%
  arrange(from,to) %>%
  mutate(from=factor(from,levels=unique(c(from,to))),
         to=factor(to,levels=levels(from)))

# for all edges
for (i in levels(files$type)) {
  write(sort(V(get(paste("tara_SUR",i,"net",sep="_")))$name),paste("tara_SUR",i,"nodes",sep="_"),ncolumns=1)
  system(paste0("sort -k 2,2 tara_SUR.suma.pairwise.invz | join -1 2 - tara_SUR_",i,
                "_nodes | sort -k 2,2 | join -1 2 - tara_SUR_",i,
                "_nodes | cut -d ' ' -f 1-2 > tara_SUR.suma.pairwise.invz.",i))
}
invz_all_taxo<-foreach(i=levels(files$type),.combine=rbind) %do% {
  read.table(paste0("tara_SUR.suma.pairwise.invz.",i),col.names=c("from","to"),stringsAsFactors=F) %>%
    mutate(type=i)
  } %>%
  left_join(tara_SUR_taxo_agg,by=c("from"="OTU")) %>%
  rename(taxo1=taxo) %>%
  left_join(tara_SUR_taxo_agg,by=c("to"="OTU")) %>%
  mutate(taxo1=ifelse(grepl("Chrysophyceae-Synurophyceae|MAST|MOCH|Bacillariophyta",taxo1),
                      sub("^(([^\\|]*\\|){4}).*$","\\1",taxo1,perl=T),
                      sub("^(([^\\|]*\\|){3}).*$","\\1",taxo1,perl=T)),
         taxo=ifelse(grepl("Chrysophyceae-Synurophyceae|MAST|MOCH|Bacillariophyta",taxo),
                     sub("^(([^\\|]*\\|){4}).*$","\\1",taxo,perl=T),
                     sub("^(([^\\|]*\\|){3}).*$","\\1",taxo,perl=T)),
         taxo1=sub("Alveolata","other_Alveolata",sub("Stramenopiles","other_Stramenopiles",sub("_X$","",sub("^.*\\|([^\\|]*)\\|$","\\1",taxo1,perl=T)))),
         taxo=sub("Alveolata","other_Alveolata",sub("Stramenopiles","other_Stramenopiles",sub("_X$","",sub("^.*\\|([^\\|]*)\\|$","\\1",taxo,perl=T))))) %>%
  group_by(type,taxo1,taxo) %>%
  summarize(n=n()) %>%
  rowwise() %>%
  mutate(from=sort(c(taxo1,taxo))[1],
         to=sort(c(taxo1,taxo))[2]) %>%
  group_by(type,from,to) %>%
  summarize(n=sum(n)) %>%
  ungroup() %>%
  arrange(from,to) %>%
  mutate(from=factor(from,levels=unique(c(from,to))),
         to=factor(to,levels=levels(from)))


# cicular summary of network eges in the inversion zone
circle_taxa<-rbind(data.frame(t="taxo_class",rbind(invz_taxo_class[,c(2,4)],setNames(invz_taxo_class[,3:4],c("from","n")))),
                   data.frame(t="all_taxo",rbind(invz_all_taxo[,c(2,4)],setNames(invz_all_taxo[,3:4],c("from","n"))))) %>%
  group_by(t,from) %>%
  summarize(n=sum(n)) %>%
  group_by(t) %>%
  mutate(percent=n/sum(n)*100) %>%
  ungroup() %>%
  filter(percent>1) %>%
  select(from) %>%
  unique() %>%
  mutate(from=sub("_"," ",from),
         color=rev(interleave(hue_pal(h=c(0,300))(length(from))))) %>%
  deframe()

for(i in c("taxo_class","all_taxo")) {
  for (j in c("cooc","coex")){
    assign(paste("invz",i,j,sep="_"),
           mutate(get(paste0("invz_",i)),from=sub("_"," ",from),to=sub("_"," ",to)) %>%
           filter(type==j,from %in% names(circle_taxa), to %in% names(circle_taxa)) %>%
             select(-type) %>%
             spread(to,n,fill=0) %>%
             column_to_rownames("from") %>%
             as.matrix())
  }
}

circos.clear()
circos.par(start.degree=90)

pdf("Figure_S8.pdf",width=9,height=9)
layout(matrix(1:4,2,2,byrow=T))
par(cex.main=1.5,cex=1)
chordDiagram(invz_all_taxo_cooc,grid.col=circle_taxa,annotationTrack="grid")
title("A",adj=0.05,line=-1)
chordDiagram(invz_all_taxo_coex,grid.col=circle_taxa,annotationTrack="grid")
title("B",adj=0.95,line=-1)
chordDiagram(invz_taxo_class_cooc,grid.col=circle_taxa,annotationTrack="grid")
title("C",adj=0.05,line=-1)
chordDiagram(invz_taxo_class_coex,grid.col=circle_taxa,annotationTrack="grid")
title("D",adj=0.95,line=-1)
leg_col<-Legend(at=c(names(circle_taxa)[15],names(circle_taxa)[-15]),type="points",
                legend_gp=gpar(col=c(circle_taxa[15],circle_taxa[-15])),pch=15,size=unit(0.8,"line"))
pushViewport(viewport(x=unit(5.05,"inches"),y=unit(4.5,"inches"), width = grobWidth(leg_col), 
                      height = grobHeight(leg_col), just = c("center","center")))
grid.draw(leg_col)
upViewport()
dev.off()

# Fold change in proportion of edges between the different clades
invz_diff<-rename(invz_all_taxo,tot=n) %>%
  right_join(invz_taxo_class) %>%
  mutate(from=sub("[-_]","\n",from),
         to=sub("[-_]","\n",to),
         type=as.factor(type),
         type=fct_relevel(type,"coex",after=1),
         type=mapvalues(type,levels(type),c("co-occurrence","co-exclusion"))) %>%
  group_by(type) %>%
  filter(from %in% sub("[-_ ]","\n",names(circle_taxa)), to %in% sub("[-_ ]","\n",names(circle_taxa))) %>%
  mutate(totp=tot/sum(tot)*100,
         np=n/sum(n)*100) %>%
  ungroup() %>%
  mutate(from=fct_rev(as.factor(from)),
         to=as.factor(to)) %>%
  rowwise() %>%
  mutate(change=ifelse(np>totp,np/totp,-totp/np))

plot_invz_heatmap<-ggplot(invz_diff,aes(to,from)) +
  geom_tile(aes(fill=change)) +
  facet_grid(~type) +
  labs(fill="fold change") +
  scale_fill_gradientn(limits=c(-12,12),values=c(0,0.5,1),
                       colors=c(colorblind_pal()(7)[7],"white",colorblind_pal()(4)[4])) +
  theme(legend.position=c(0.6,0.25),
        axis.title=element_blank(),
        axis.text.x=element_text(angle=90,hjust=1,vjust=0.5))
ggsave("Figure_4.pdf",plot_invz_heatmap,width=9,height=5)


# for DCM
invz_taxo_class_DCM<-filter(invz_taxo,project=="tara_DCM") %>%
  select(-project,-from,-to) %>%
  mutate(taxo1=ifelse(grepl("Chrysophyceae-Synurophyceae|MAST|MOCH|Bacillariophyta",taxo1),
                      sub("^(([^\\|]*\\|){4}).*$","\\1",taxo1,perl=T),
                      sub("^(([^\\|]*\\|){3}).*$","\\1",taxo1,perl=T)),
         taxo=ifelse(grepl("Chrysophyceae-Synurophyceae|MAST|MOCH|Bacillariophyta",taxo),
                     sub("^(([^\\|]*\\|){4}).*$","\\1",taxo,perl=T),
                     sub("^(([^\\|]*\\|){3}).*$","\\1",taxo,perl=T)),
         taxo1=sub("Alveolata","other_Alveolata",sub("Stramenopiles","other_Stramenopiles",sub("_X$","",sub("^.*\\|([^\\|]*)\\|$","\\1",taxo1,perl=T)))),
         taxo=sub("Alveolata","other_Alveolata",sub("Stramenopiles","other_Stramenopiles",sub("_X$","",sub("^.*\\|([^\\|]*)\\|$","\\1",taxo,perl=T))))) %>%
  rowwise() %>%
  mutate(from=sort(c(taxo1,taxo))[1],
         to=sort(c(taxo1,taxo))[2]) %>%
  group_by(type,from,to) %>%
  summarize(n=n()) %>%
  ungroup() %>%
  arrange(from,to)

# for all edges
for (i in levels(files$type)) {
  write(sort(V(get(paste("tara_DCM",i,"net",sep="_")))$name),paste("tara_DCM",i,"nodes",sep="_"),ncolumns=1)
  system(paste0("sort -k 2,2 tara_DCM.suma.pairwise.invz | join -1 2 - tara_DCM_",i,
                "_nodes | sort -k 2,2 | join -1 2 - tara_DCM_",i,
                "_nodes | cut -d ' ' -f 1-2 > tara_DCM.suma.pairwise.invz.",i))
}
invz_all_taxo_DCM<-foreach(i=levels(files$type),.combine=rbind) %do% {
  read.table(paste0("tara_DCM.suma.pairwise.invz.",i),col.names=c("from","to"),stringsAsFactors=F) %>%
    mutate(type=i)
} %>%
  left_join(tara_DCM_taxo_agg,by=c("from"="OTU")) %>%
  rename(taxo1=taxo) %>%
  left_join(tara_SUR_taxo_agg,by=c("to"="OTU")) %>%
  mutate(taxo1=ifelse(grepl("Chrysophyceae-Synurophyceae|MAST|MOCH|Bacillariophyta",taxo1),
                      sub("^(([^\\|]*\\|){4}).*$","\\1",taxo1,perl=T),
                      sub("^(([^\\|]*\\|){3}).*$","\\1",taxo1,perl=T)),
         taxo=ifelse(grepl("Chrysophyceae-Synurophyceae|MAST|MOCH|Bacillariophyta",taxo),
                     sub("^(([^\\|]*\\|){4}).*$","\\1",taxo,perl=T),
                     sub("^(([^\\|]*\\|){3}).*$","\\1",taxo,perl=T)),
         taxo1=sub("Alveolata","other_Alveolata",sub("Stramenopiles","other_Stramenopiles",sub("_X$","",sub("^.*\\|([^\\|]*)\\|$","\\1",taxo1,perl=T)))),
         taxo=sub("Alveolata","other_Alveolata",sub("Stramenopiles","other_Stramenopiles",sub("_X$","",sub("^.*\\|([^\\|]*)\\|$","\\1",taxo,perl=T))))) %>%
  group_by(type,taxo1,taxo) %>%
  summarize(n=n()) %>%
  rowwise() %>%
  mutate(from=sort(c(taxo1,taxo))[1],
         to=sort(c(taxo1,taxo))[2]) %>%
  group_by(type,from,to) %>%
  summarize(n=sum(n)) %>%
  ungroup() %>%
  arrange(from,to)

# cicular summary of network eges in the inversion zone
circle_taxa_DCM<-rbind.data.frame(data.frame(t="taxo_class",rbind(invz_taxo_class_DCM[,c(2,4)],setNames(invz_taxo_class_DCM[,3:4],c("from","n")))),
                                  data.frame(t="all_taxo",rbind(invz_all_taxo_DCM[,c(2,4)],setNames(invz_all_taxo_DCM[,3:4],c("from","n"))))) %>%
  group_by(t,from) %>%
  summarize(n=sum(n)) %>%
  group_by(t) %>%
  mutate(percent=n/sum(n)*100) %>%
  ungroup() %>%
  filter(percent>1) %>%
  select(from) %>%
  unique() %>%
  mutate(from=sub("[-_]","\n",from),
         color=rev(interleave(hue_pal(h=c(0,300))(length(from))))) %>%
  deframe()

# Fold change in proportion of edges between the different clades
invz_diff_DCM<-rename(invz_all_taxo_DCM,tot=n) %>%
  right_join(invz_taxo_class_DCM) %>%
  mutate(from=sub("[-_]","\n",from),
         to=sub("[-_]","\n",to),
         type=as.factor(type),
         type=fct_relevel(type,"coex",after=1),
         type=mapvalues(type,levels(type),c("co-occurrence","co-exclusion"))) %>%
  group_by(type) %>%
  filter(from %in% names(circle_taxa_DCM), to %in% names(circle_taxa_DCM)) %>%
  mutate(totp=tot/sum(tot)*100,
         np=n/sum(n)*100) %>%
  ungroup() %>%
  mutate(from=fct_rev(as.factor(from)),
         to=as.factor(to)) %>%
  rowwise() %>%
  mutate(change=ifelse(np>totp,np/totp,-totp/np))

plot_invz_heatmap_DCM<-ggplot(invz_diff_DCM,aes(to,from)) +
  geom_tile(aes(fill=change)) +
  facet_grid(~type) +
  labs(fill="fold change") +
  scale_fill_gradientn(limits=c(-12,12),values=c(0,0.5,1),
                       colors=c(colorblind_pal()(7)[7],"white",colorblind_pal()(4)[4])) +
  theme(legend.position=c(0.6,0.25),
        axis.title=element_blank(),
        axis.text.x=element_text(angle=90,hjust=1,vjust=0.5))
ggsave("Figure_S9.pdf",plot_invz_heatmap_DCM,width=9,height=5)
```

### 8 Dependencies

#### 8.1 bash

```
# GNU bash (4.4.12(1))
# GNU parallel (20190822)
# SUMATRA (1.0.34)
# MAFFT (7.407)
# Open MPI (2.0.2)
# RAxML (8.2.12)
# slurm (16.05.9)
```

#### 8.2 R session informations

```
R version 3.5.2 (2018-12-20)
Platform: x86_64-pc-linux-gnu (64-bit)
Running under: Ubuntu 18.04.4 LTS

Matrix products: default
BLAS: /usr/lib/x86_64-linux-gnu/blas/libblas.so.3.7.1
LAPACK: /usr/lib/x86_64-linux-gnu/lapack/liblapack.so.3.7.1

locale:
 [1] LC_CTYPE=en_US.UTF-8       LC_NUMERIC=C               LC_TIME=de_CH.UTF-8       
 [4] LC_COLLATE=en_US.UTF-8     LC_MONETARY=de_CH.UTF-8    LC_MESSAGES=en_US.UTF-8   
 [7] LC_PAPER=de_CH.UTF-8       LC_NAME=C                  LC_ADDRESS=C              
[10] LC_TELEPHONE=C             LC_MEASUREMENT=de_CH.UTF-8 LC_IDENTIFICATION=C       

attached base packages:
[1] parallel  grid      stats     graphics  grDevices utils     datasets  methods   base     

other attached packages:
 [1] knitr_1.23            abind_1.4-5           rhdf5_2.26.2          doParallel_1.0.14    
 [5] picante_1.7           nlme_3.1-137          biomformat_1.10.1     Hmisc_4.2-0          
 [9] Formula_1.2-3         survival_2.43-3       vegan_2.5-4           lattice_0.20-38      
[13] permute_0.9-4         iterators_1.0.10      foreach_1.4.4         forcats_0.4.0        
[17] stringr_1.4.0         dplyr_0.8.3           purrr_0.3.1           readr_1.3.1          
[21] tidyr_1.0.0           tibble_2.1.3          tidyverse_1.2.1       plyr_1.8.4           
[25] ggthemes_4.1.1        scales_1.0.0          ggnetwork_0.5.8       ggpubr_0.2.5         
[29] magrittr_1.5          ggplot2_3.1.0         igraph_1.2.4          ComplexHeatmap_1.20.0
[33] circlize_0.4.8        ape_5.2              

loaded via a namespace (and not attached):
 [1] lubridate_1.7.4     RColorBrewer_1.1-2  httr_1.4.0          tools_3.5.2        
 [5] backports_1.1.3     utf8_1.1.4          R6_2.4.0            rpart_4.1-13       
 [9] lazyeval_0.2.1      mgcv_1.8-27         colorspace_1.4-0    nnet_7.3-12        
[13] GetoptLong_0.1.8    withr_2.1.2         tidyselect_0.2.5    gridExtra_2.3      
[17] compiler_3.5.2      cli_1.0.1           rvest_0.3.4         htmlTable_1.13.1   
[21] xml2_1.2.0          labeling_0.3        checkmate_1.9.1     digest_0.6.18      
[25] foreign_0.8-70      rmarkdown_1.13      base64enc_0.1-3     pkgconfig_2.0.2    
[29] htmltools_0.4.0     htmlwidgets_1.3     rlang_0.4.0         GlobalOptions_0.1.1
[33] readxl_1.3.1        rstudioapi_0.9.0    shape_1.4.4         generics_0.0.2     
[37] jsonlite_1.6        acepack_1.4.1       Matrix_1.2-15       Rhdf5lib_1.4.2     
[41] Rcpp_1.0.3          munsell_0.5.0       fansi_0.4.0         lifecycle_0.1.0    
[45] stringi_1.3.1       yaml_2.2.0          MASS_7.3-51.1       crayon_1.3.4       
[49] haven_2.1.0         cowplot_0.9.4       splines_3.5.2       hms_0.4.2          
[53] zeallot_0.1.0       pillar_1.3.1        rjson_0.2.20        ggsignif_0.6.0     
[57] reshape2_1.4.3      codetools_0.2-16    glue_1.3.0          evaluate_0.13      
[61] latticeExtra_0.6-28 data.table_1.12.0   modelr_0.1.4        vctrs_0.2.0        
[65] cellranger_1.1.0    gtable_0.2.0        assertthat_0.2.0    xfun_0.5           
[69] broom_0.5.2         cluster_2.0.7-1     ellipsis_0.3.0
```
